# Representation in genetic studies affects inference about genetic architecture

**DOI:** 10.64898/2026.01.12.699135

**Authors:** Jared M. Cole, Shane Rybacki, Samuel Pattillo Smith, Olivia S. Smith, Arbel Harpak

**Affiliations:** Department of Integrative Biology, University of Texas at Austin, Austin, TX, USA; Department of Population Health, University of Texas at Austin, Austin, TX, USA

## Abstract

Knowledge of a trait’s “genetic architecture,” namely the joint distribution of allele frequencies of causal variants and the direction and magnitude of their effects, is essential to understanding its evolution and underlying biology. Inferences about genetic architecture are based on data collected in different ways in cohorts recruited through heterogeneous mechanisms. As a result, cohorts differ in genotype, environment, and trait distributions. For example, the UK Biobank (UKB) was designed for broad population representation, whereas FinnGen drew extensively from clinical registries enriched for diagnosed health conditions. Here, we asked whether representation in genetic studies influences inferences about genetic architectures. Using GWAS data from the UKB, All of Us (AoU), and FinnGen, we compared several summaries of genetic architecture. We find systematic differences in some summaries, such as SNP heritability, which were often (in 6/13 traits) significantly lower in AoU than in the UKB (and never the reverse), even when matching samples such that they have similar genetic ancestry compositions. This result aligns with other recent evidence that biobanks enriched for diagnosed health conditions, also sometimes characterized by less-standardized phenotyping, have lower heritability than population-based biobanks. We highlight a second case, where a summary of genetic architecture varies considerably but not systematically across traits and biobanks. Such is the case for the mean direction of allelic effects (“sign bias”). For example, 72% of rare minor alleles affecting type 2 diabetes risk are inferred to be risk-increasing based on AoU data, while nearly all (*>*99%) are inferred to be risk-increasing based on UKB data. We hypothesize that the inferred sign bias is heavily influenced by the skewness of the trait distribution in the study and otherwise largely independent of other study or trait characteristics, including whether the trait is binary or quantitative. We provide strong support for this hypothesis through simulations and data from the three biobanks: the variation in inferred sign bias for rare minor alleles across traits and biobanks is explained remarkably well (82% and 97% of variance explained for trait-associated SNPs and randomly selected SNPs, respectively) solely by the trait’s skewness in the biobank, with residual biobank-specificity explaining little. Our findings suggest that inferences about the map between genetic and trait variation can depend on study design and participation in genetic studies in surprising ways.

## Introduction

Characterizing the mapping of genetic variation onto trait variation is central to understanding the biological mechanisms underlying complex traits and their evolution. Summaries of this mapping for a given trait are often referred to jointly as its “genetic architecture.” More formally, the genetic architecture of a trait comprises the joint distribution of frequencies and effects (including both magnitudes and directions) of causal variants^1,2^. Genome-wide association studies (GWAS) have allowed researchers to estimate variant-trait associations for hundreds of complex traits and diseases and thus characterize several aspects of their underlying genetic architectures^3^.

Through this effort, we have learned that many traits are highly polygenic, i.e. most trait variation is explained by small effects of numerous alleles^1,4–7^. Allelic effects typically span multiple orders of magnitude (such that a few large effects are present amid many tiny ones^8,9^) and variants tend to exhibit substantial pleiotropy, influencing multiple traits simultaneously^10,11^. Genetic architecture is further molded by evolutionary forces such as mutation, natural selection, and genetic drift, which produce recognizable relationships between effect magnitude and allele frequency. One famous example is the negative monotonic relationship between the average magnitude of allelic effects and their frequency^1,12,13^. Since inferences about genetic architecture are key to our understanding of the evolution and biology of complex traits, and because they are often assumed to be intrinsic to traits, it is concerning when aspects of architecture are found to vary across studies^14–17^. This variation is often attributed primarily to variation in genetic ancestry across study cohorts, leading to variable genetic interactions^18–21^. However, studies also differ by contexts, such as country, healthcare systems, or recruitment strategies. As one example, a biobank that primarily recruits from specialty clinics will over-represent individuals with severe or multiple diseases (e.g., FinnGen), whereas a population-volunteer cohort (e.g., UK Biobank) will under-represent people who are socioeconomically disadvantaged or in poor health. Differences in cohort recruitment can influence cohort-level estimates of allelic effects themselves and subsequent summaries such as genetic correlations^22–24^. There are existing approaches that statistically adjust for heterogeneity across studies (e.g., inverse-probability weighting^25,26^), but those do not directly inquire about its drivers. Thus, it remains unclear how study cohort characteristics influence inferences about the genetic architecture of complex traits.

Here, we ask how representation in genetic studies affects inferences about genetic architecture. We study differences in summaries of genetic architecture across three major biobanks with distinct study design choices: a population-volunteer cohort (UK Biobank^27^), a cohort emphasizing inclusion of under-represented groups (All of Us^28^), and a diagnosis-enriched cohort (FinnGen^29^). We compare summaries across 14 traits spanning both quantitative traits and disease endpoints. We then focus on the mean direction of allelic effects (“sign bias”) as a tractable case study. Sign bias estimates vary considerably across biobanks for some traits. We hypothesize that trait skewness may affect virtually any method used to estimate sign bias, and show support for this hypothesis both with empirical analysis across biobanks and traits and in simulation studies under a null scenario of no true sign bias. Together, these findings show that our estimates of key summaries of genetic architecture can depend heavily on properties of study design, including how participants are recruited and how traits are measured.

## Results

### Headline summaries of genetic architecture vary across biobanks

We investigated whether inferences about genetic architectures could depend on characteristics of recruitment and measurement in genetic studies. To do so, we compared summaries of genetic architecture as estimated using GWAS data from distinct biobanks.

The UK Biobank (UKB^30^) is a population-based cohort in the United Kingdom that was recruited from over half a million adults aged 40–69 from 2006–2010 located near assessment centers, where a large collection of traits were measured and biological samples taken at enrollment. Follow-up assessments and monitoring of health records provided additional measurements. Subsequent studies have found that the selective participation into the cohort has led to a pronounced healthy-volunteer bias^31^. All of Us (AoU^28^) represents a broader, longitudinal cohort designed to raise participation among groups historically underrepresented in biomedical research. It is an ongoing study that aims to recruit a minimum of 1 million participants that are representative of the US population. Participants are recruited through various means, such as through direct enrollment, healthcare organizations, and community-based efforts. Traits are measured using various approaches, largely via electronic health records, and participants were not generally recruited on the basis of specific diagnoses. Nevertheless, due to the heavily healthcare-based recruitment, AoU is significantly more enriched for diseases and clinical conditions relative to UKB^32^.

We analyzed GWAS summary statistics for 14 traits from the UKB^27^ and the AoU databases using ancestry-matched samples across a common set of approximately 10 million SNPs (see **Methods**). We focused on three morphological traits, six hematological traits (four of which are immune-related white blood cell traits and two red blood cell traits), and five disease traits (**Fig. S1**; **Table S1**). All traits were analyzed in their raw units, with no standardization or other transformation (see our reasoning for this choice in the **Discussion**). Using bivariate LD Score Regression (LDSC^33^), we estimated SNP heritability (SNP *h*^2^) for each trait-biobank pair and calculated their cross-biobank genetic correlations. We also estimated the effective polygenicity of each trait. Effective polygenicity is a measure of how evenly the heritable signal is distributed across variants. If the majority of the trait’s heritability is due to only a few loci, then the effective polygenicity is small, but if it is due to many loci across the genome, the effective polygenicity is large^34,35^. We note that this measure differs from commonly used measures of polygenicity that probe the number of variants with non-zero effects^35–38^.

While the subsets of UKB and AoU we used for GWAS were, by design, similar in their genetic ancestry composition, summaries of genetic architecture often differed. In particular, SNP heritability estimates in AoU were lower than those in UKB (approximately 17% ± 4% lower on average across traits, estimated using Deming regression; **Fig. 1A**; **Table S2**). For binary diseases, we estimated SNP heritability on the liability-scale^39^ (observed-scale estimates shown in **Fig. S10**), treating the disease prevalence within each biobank as identical to the population prevalence. We made this choice because the cohorts are sampled in different countries and contexts, and, after subsetting and filtering, estimating the prevalence in an orthogonal way (other than based on the sample prevalence) is nontrivial. We note that if we exclude the binary traits altogether, the estimates in AoU appear even lower than their UKB equivalents (approximately 18% ± 4% lower on average). These results are consistent with prior reports of lower SNP heritability in AoU than UKB, even when GWAS in the two studies showed high genetic correlations^40^.

**Figure 1:**
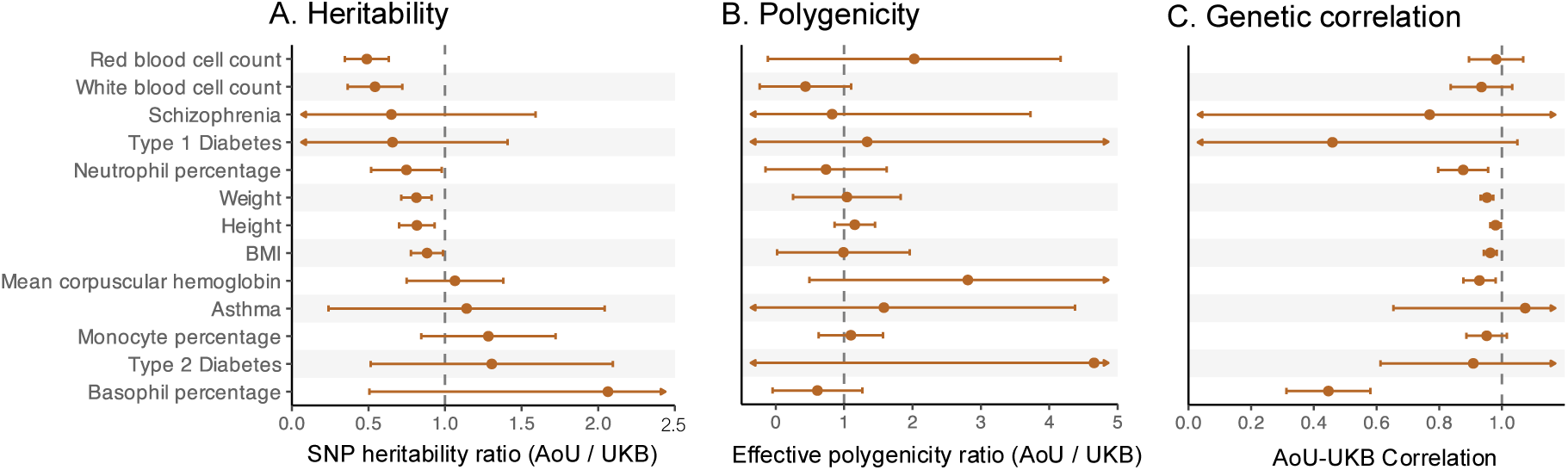
Summaries of genetic architecture vary across biobanks. For each trait (row), shown is the correspondence of estimates derived from GWAS performed in the All of Us (AoU) and UK Biobank (UKB) datasets. (A) The ratio of the SNP heritability estimated using LD Score Regression (with liability scale estimates for binary traits; observed scale estimates shown in **Fig. S10**and reported in **Table S2**). Across traits, AoU estimates of SNP heritability are on average lower than UKB. (B) Ratio of effective polygenicity, which summarizes how diffuse vs. concentrated a trait’s heritability is across genomic loci. (C) Each row shows the genetic correlation, estimated using bivariate LD Score Regression, between allelic effects (on the same trait) estimated in UKB and AoU GWAS. Points represent estimates and horizontal bars represent 95% confidence intervals. Confidence intervals that extend beyond the displayed axis ranges are truncated, with arrows indicating the direction where the interval continues. Dashed vertical lines correspond to a ratio of 1 in A and B and a perfect genetic correlation (*r_g_* = 1) in C. Alzheimer’s disease (AD) is omitted from the above panels because the SNP heritability estimates in UKB were negative, precluding valid comparisons of polygenicity and genetic correlations.

While heritability is a major focus in studies of genetic architecture, it is not an immutable property of a trait and should not be expected to be the same in distinct samples, even with close ancestry matching^22,41–43^. Here, the GWAS samples we analyzed are from different countries and societies. Further, they came about through fundamentally different recruitment strategies and different methods of data collection. These features of a study carry several implications for heritability. For example, the composition of (largely unmeasured) environmental and social factors is plausibly different across samples, resulting in differences in the extent of trait variation that is attributable to genetic variation.

Nevertheless, given that the statistical power to detect genotype-trait associations in biomedical GWAS directly depends on heritability^44–46^, it is noteworthy that heritability is lower in AoU for the biomedical traits we examined. These results add to recent observations^16,40^ that together support a surprising conclusion: across four biobanks, it appears that SNP heritability tends to be lower in biobanks with disease-enriched recruitment. To our knowledge, our analysis is the first to demonstrate this difference based on an ancestry-matched comparison. In the **Discussion**, we suggest a few hypotheses for the causes of this difference.

Other quantities, like the dispersion of signal across variants (polygenicity) or the correlation of causal allelic effects (genetic correlation) among the samples are thought to be closer to trait-level properties because they do not directly depend on environmental variance and because causal genetic effects are expected to be similar across populations^18,19,35,44,47–49^. For most traits, effective polygenicity indeed differed only slightly between biobanks (**Fig. 1B**; **Table S3**), with type 2 diabetes and mean corpuscular hemoglobin (AoU/UKB ratio of 4.65 ± 3.09 and 2.81 ± 1.18, respectively) showing the largest (though statistically insignificant) differences. Though genetic correlations are in general high (0.86 on average) and are consistent with previously reported estimates between primarily European ancestry subsamples of UKB and AoU^40^, our results identify significant departures from perfect correlations for 6 out of the 13 traits examined (**Fig. 1C**; **Table S4**). The most notable examples are basophil percentage (AoU-UKB genetic correlation of 0.45±0.07) and neutrophil percentage (AoU-UKB genetic correlation of 0.88±0.04). To confirm that these deviations from a genetic correlation of 1 are not a result of sampling noise, we estimated genetic correlations for random subsets of the UKB. Across traits, these never significantly deviated from one (**Text S1**; **Table S4**; **Fig. S9**). This finding is consistent with recent work showing that estimates of genetic correlations can depend on study design. Song et al.^23^ demonstrated that heritable participation biases, for example, can impact estimates of cross-trait genetic covariance.

### Sign bias: A case study of drivers of cross-biobank differences in genetic architecture

The comparisons of SNP heritability, polygenicity and causal effects (probed through genetic correlations) may together suggest that genetic architectures vary across biobanks. To investigate the idea that this is largely a result of characteristics of the study sample, we examined sign bias, which is a specific, tractable summary of allelic effects on complex traits.

We defined sign bias for a given set of variants and traits as the mean direction (sign) of allelic effects on the trait among these variants. The direction ought to be defined with respect to some well-defined reference allele across variants. Here, we used the sign of the minor allele throughout. Hypothetically, sign bias may reflect biological mechanisms that couple molecular traits (e.g., gene expression levels) to downstream effects on complex traits, or it may reflect the mode of natural selection acting on those traits^1,50–52^. For example, traits under directional selection favoring an increase in population mean will accumulate more trait-decreasing minor alleles than trait-increasing ones, while those under stabilizing selection will show no sign bias^13,53–56^. At the same time, sign bias should be more tractable and less noisily estimated than the full distribution of allelic effects^57^.

In the **Methods** section, we describe a procedure that we developed to estimate sign bias for a set of variants. In short, we applied adaptive shrinkage (ash^58^), an empirical Bayes method that incorporates uncertainty in allelic effects, to sets of variants accounting for linkage disequilibrium. We estimated sign bias in sets of the most strongly associated SNPs across traits, bins of minor allele frequency and biobanks (**Table S5**). In what follows, we further focus on the most significantly associated SNPs for each trait (in **Figs. S3**, **S5**, and **S7**, we present results obtained with SNPs sampled randomly with respect to their trait association). Finally, to expand our scope, we also analyzed data from a third biobank, FinnGen^29^, for five corresponding traits also present in UKB and AoU for which the necessary summary data were publicly available. FinnGen is a nationwide study in Finland that recruited approximately 500,000 people in 2023. The cohort was deliberately enriched for diagnosed diseases by recruiting primarily through digital health registries and hospital clinics. We note that for these FinnGen traits, the GWAS were performed by other researchers and did not include any ancestry matching to the UKB and AoU subsets that we analyzed.

**Figure 2:**
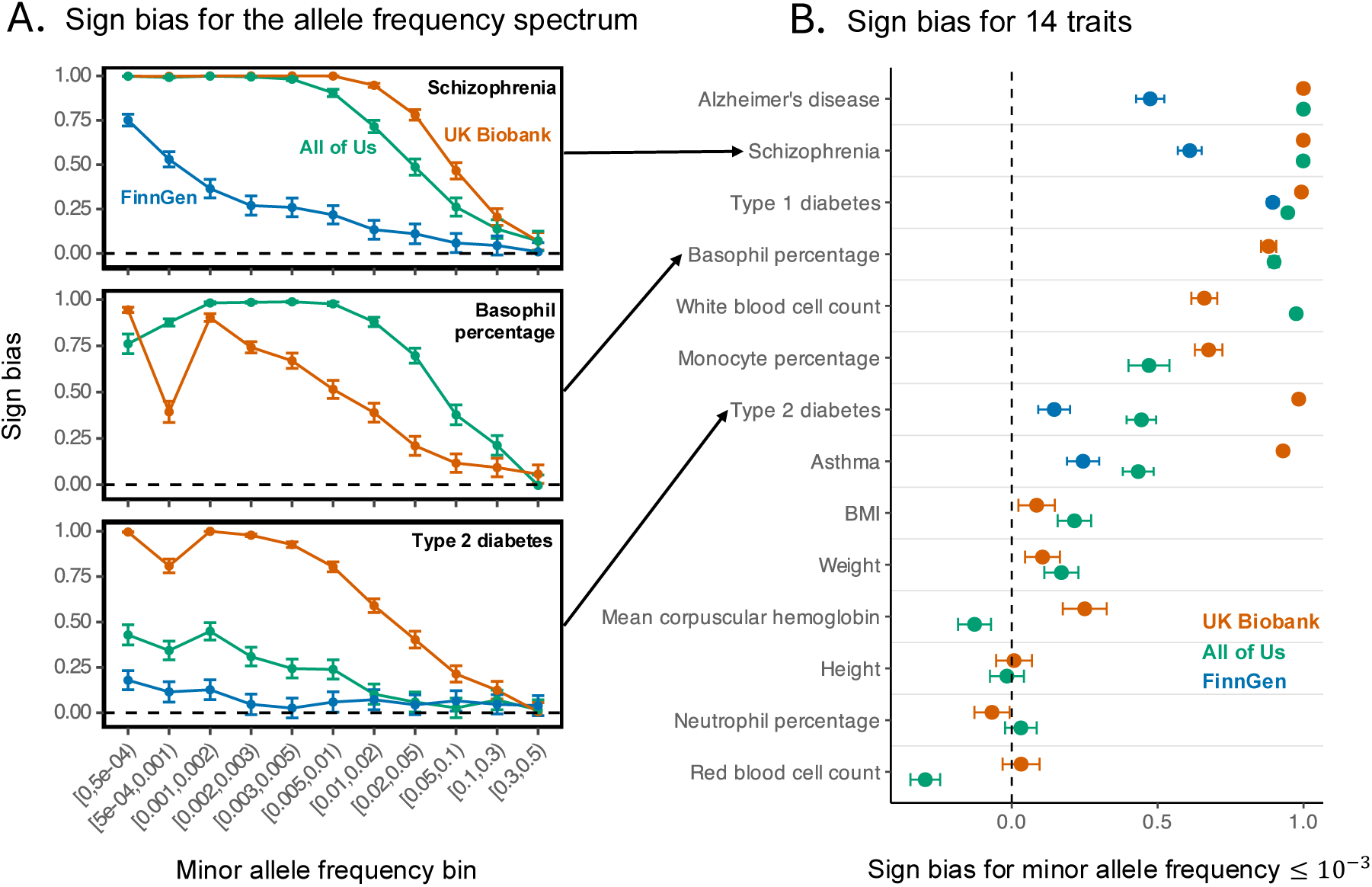
Sign bias varies across biobanks. We investigated the mean sign of allelic effects (“sign bias”) as an example summary of genetic architecture that can depend on the study cohort in which it is estimated. For each trait and a range of minor allele frequencies, we considered the sign of one SNP per approximately independent LD block. Here, we specifically used SNPs that were the most significantly associated with the trait in the block (**Figs. S3** and **S5** show a reproduction of the analysis with randomly selected SNPs, i.e. regardless of the significance of trait association). The sign bias for each set of SNPs ranges from -1 (all SNPs are trait/risk decreasing) to 1 (all SNPs are trait/risk increasing). The sign bias is estimated using empirical Bayes adaptive shrinkage (ash) to account for measurement uncertainty. Points show estimates and error bars show the 95% confidence intervals. Dashed horizontal and vertical lines show no sign bias (i.e., a value of 0). (A) Rare alleles are more sign-biased. Shown are three representative traits, with other traits shown in **Fig. S2**. (B) The estimated sign bias can vary substantially across three biobanks. Focusing on the rarer alleles (minor allele frequency of up to 0.1%, other frequency thresholds are shown in **Fig. S4**), diseases tended to show the strongest positive sign bias, whereas morphometric traits (height, weight, BMI) showed little bias. Estimates from All of Us, the UK Biobank and FinnGen were highly discrepant for many of the traits analyzed.

**Figure 3:**
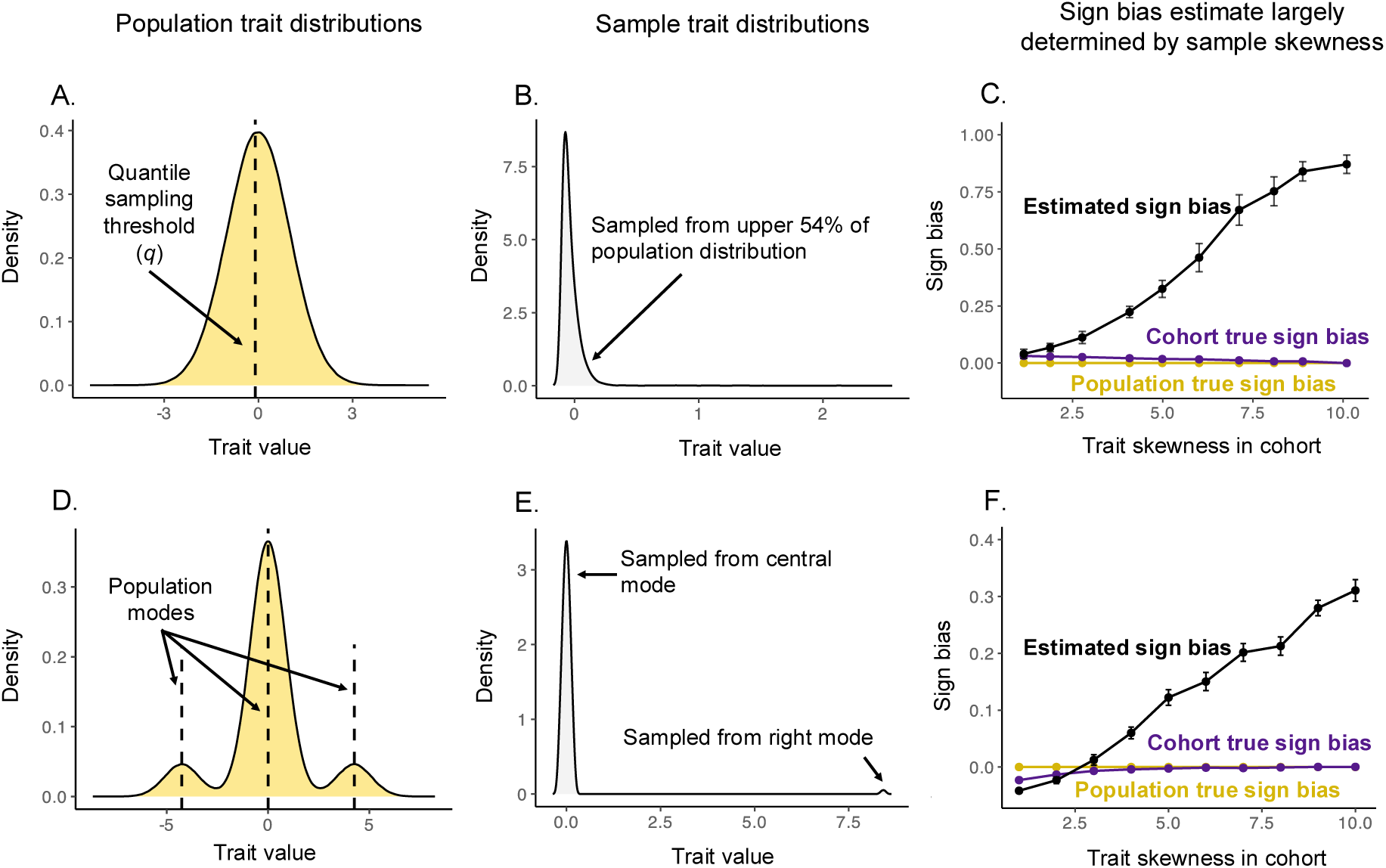
Simulations illustrating that skewness of the cohort trait distribution can translate to sign bias. We used two distinct population distributions with corresponding distinct non-random sampling schemes to generate cohorts of variable trait skewness. Upper row: **(A)** We simulated a population of 2 *×* 10^6^ individuals with equal numbers of trait-increasing and trait-decreasing minor alleles of equal magnitude effects and matched allele frequencies. **(B)** We drew a non-random sample of 10,000 individuals by sampling only above a quantile cutoff (*q*) of the population trait distribution. Shown here is an example with *q* = 0.46, corresponding to a trait value threshold of *−*0.1 and a skew of *≈* 10. **(C)** Each data point shows the average sign bias (with bars showing 95% CIs) for minor alleles with frequencies up to 1% across 20 iterations for a specific value of *q* and the corresponding mean realized trait skewness across iterations. The population sign bias is zero by construction (yellow). Sampling only induces a small sign bias (measured as the average sign of minor alleles segregating in the cohort, dark purple)—but the asymmetric effect of cohort skewness on trait-increasing and trait-decreasing minor alleles leads to a pronounced, monotonic relationship between cohort skewness and inferred sign bias (black). Lower row: **(D)** We simulated a population as in the top row, but whose trait distribution is a mixture of three latent modes (yellow; dashed lines mark mode centers). **(E)** We drew a non-random sample of 10,000 individuals as a mixture of two components: a large fraction sampled, with low variance, from the central mode, and a small fraction from the right mode. Varying the fraction drawn from the right mode modulates cohort skewness. An example right-skewed cohort (skew *≈* 10; gray) is shown by sampling 0.9% of individuals from the right mode. **(F)** Same as in panel C, but results are shown for the tri-modal sampling scheme illustrated in D and E.

Across traits, sign bias was negligible at common alleles (minor allele frequencies ≥ 0.1) but increased for rarer alleles (**Figs. 2A, S2**). In what follows, we therefore focus on the latter (but see **Fig. S4** for complementary results including more common alleles). While allelic effect estimates for rare alleles are often noisily estimated^59^, our sign bias estimation method is designed to capture the mean sign accurately, and weighs evidence across variants by confidence in their sign (**Methods**).

The bias was most pronounced for binary disease traits, such as schizophrenia, type 1 diabetes, and Alzheimer’s disease (towards risk-increasing minor alleles) (**Fig. 2B**). In contrast, morphological traits (e.g., height, weight, BMI) showed little to no bias. However, it was evident that sign bias could not be viewed as a strictly trait-level property, as estimates can differ dramatically across biobanks. For example, among the most significantly trait-associated minor alleles with frequencies up to 0.1% (shown in **Fig. 2B**), approximately 99.2% of variants for type 2 diabetes were inferred to be risk-increasing in UKB, but only 72.2% in AoU and 57.3% in FinnGen. Asthma was similarly discrepant, with 96.5% inferred to be risk-increasing in UKB, compared to 71.7% for AoU and 62.2% for FinnGen. By contrast, schizophrenia and Alzheimer’s disease were nearly saturated in UKB and AoU (*>*99% of minor alleles inferred to be risk-increasing), whereas FinnGen estimates were substantially lower (85.5% for schizophrenia and 73.7% for Alzheimer’s disease). Mean corpuscular hemoglobin showed opposite patterns between the UKB and AoU: in the UKB, the inferred sign bias was positive (62.5% of variants were estimated to increase the trait), whereas in AoU it was slightly negative (only 43.6% were estimated to increase the trait).

That sign biases vary across biobanks indicates that they do not reliably reflect the impact of de novo mutations on the trait. Similarly, sign biases are unlikely to reliably reflect the mode or intensity of natural selection, especially given that two of the three GWAS samples were ancestry-matched. What, then, could explain the marked variation in sign bias across biobanks?

### Skewness of the trait distribution in a study strongly predicts sign bias

We hypothesize that differences in estimated sign bias track differences in trait distribution across biobank cohorts, and specifically their skewnesses (standardized third central moments). Our intuition stems from two aspects of association asymmetry (favoring one sign) for binary traits—discoveries and spurious associations—for skewed trait distributions.

First, we considered discovery of risk vs. protective variants in binary disease traits. When cases are rare, observing even a few carriers of the same allele among cases hints at the allele being risk-increasing. On the other hand, the identification of a protective effect of the same magnitude would require observing carriers of the allele who would have otherwise been likely cases (i.e., their background risk, excluding the focal variant, is high) but remain unaffected. Previous work showed that the strength of association between allele and disease status, and resultant test statistics, are indeed bounded by a function of the marginal frequency of the allele and the incidence of the disease^60–62^. When the minor allele increases risk for a rare disease, carriers can be concentrated among the relatively few cases, allowing a large (positive) correlation; yet if the minor allele is protective with the same magnitude of effect, its frequency among non-cases cannot be as high and its (negative) correlation with disease is constrained to be weaker.

The implications extend beyond the power asymmetry for large effect alleles and extend to quantitative traits, particularly to a discrepancy in the uncertainty of allelic effect estimates between trait-increasing and trait-decreasing alleles. If the trait distribution in the cohort is positively skewed, a minor allele can more strongly correlate with the trait if it is trait-increasing and enriched in the right tail. If, in contrast, a minor allele is trait-decreasing and depleted in the right tail, its correlation with the trait would be limited. This can be viewed generally as a coupling problem: given fixed marginals for a rare genotype and a skewed trait, only certain dependence structures are possible such that the maximum correlation depends on how well the skewness of the two variables can be aligned^62^.

Second, we considered previous work on false discovery asymmetry in skewed distributions, which also becomes more pronounced for rare alleles^63,64^. In sum, if the trait distribution is positively skewed in a sample, the correlation discrepancy should translate to higher uncertainty in allelic effect estimation for trait-decreasing minor alleles. This could impact many downstream inferences about genetic architecture, including the sign bias.

Using a simulation study (**Methods**), we confirmed that participation-induced trait skewness can influence estimation of sign bias. In short, we studied two cohort sampling designs (**Fig. 3**). In both cases, the underlying population contains equal numbers of trait-increasing and trait-decreasing variants with equal magnitudes of effects. In the first scenario, the trait distribution in the population is Normal (**Fig. 3A**). We generated right-skewed cohorts by preferentially sampling individuals above a threshold. Here, the threshold determined the skewness in the cohort (**Fig. 3B**). In the second design, we generated a population with a tri-modal, symmetric trait distribution (**Fig. 3D**). We then sampled a cohort only from the central and the right modes. Here, the fraction drawn from the right mode determined cohort skewness (**Fig. 3E**). We note that the “true” sign bias in the cohort (defined by the direction of effects among variants segregating in the cohort) changed little with skewness (dark purple points), but in a different fashion in the two scenarios (dark purple lines in **Fig. 3C,F**). In both scenarios, the estimated sign bias increases monotonically with skewness (black points in **Fig. 3C,F, Table S6**).

While the simulations in **Fig. 3** demonstrate that skewed cohort sampling can induce sign bias, they cannot capture the extent to which the pattern is due to an asymmetry in the ability to detect true trait-increasing vs trait-decreasing alleles and false-positives due to spurious associations, as discussed above. We therefore repeated the first simulation under a scenario where all alleles have zero effect on the trait. The estimated sign bias again increased with cohort skewness, even with no true sign bias in the population (**Fig. S11**). While both mechanisms likely contribute, these results show that false positive associations are sufficient on their own to generate the observed qualitative pattern. Inverse-rank normalization of the sampled cohort trait distribution largely attenuated this relationship, indicating that the relationship is driven by the skewness of the distribution in the cohort (**Fig. S11**).

The expectation that trait skewness alone should predict inferred sign biases also held in empirical biobank data. Because the three biobank samples are from different countries and the study designs have somewhat different goals and recruitment approaches, skewness varied substantially for some traits. For example, the number of schizophrenia cases varied (translating to a variable skew) across biobanks (a sample prevalence of 0.05% in UKB as compared to 0.5% in AoU); similarly, white blood cell count, a quantitative trait, was markedly more skewed in AoU (42.7) than in UKB (10.6) in the ancestry-matched samples we considered. These differences likely reflect, in part, differences among biobanks in both recruitment and measurement (**Discussion**). For example, for a binary disease, greater recruitment of cases makes the case/non-case distribution less imbalanced and therefore less skewed. Consequently, diagnosis-enriched studies like FinnGen will tend to be less skewed than population-based cohorts where cases are rarer.

We focused again on rare minor alleles for the most trait-associated variants (minor allele frequencies ≤ 0.1% in **Fig. 2B**). The sign bias for diseases with substantial case imbalance (Alzheimer’s disease, schizophrenia, and type 1 diabetes) or quantitative traits with extreme skew (like white blood cell count in AoU) was nearly one, indicating the vast majority of rare minor alleles were risk-or trait-increasing. The sign bias of traits with a more moderate skew, like type 2 diabetes and basophil percentage, was smaller but also substantial. Morphometric traits with vanishingly little skewness showed no sign bias (**Fig. 4A; Table S7**). Across all traits, sign bias and trait skewness are highly correlated irrespective of the biobank (pooled Spearman’s *ρ* = 0.967, *P <* 0.001; **Table S8**). A quadratic logit fit supported a saturating, curved relationship of sign bias with skewness (adjusted *R*^2^ = 0.82, *P <* 0.001 after removing one outlier, white blood cell count, with simultaneously extreme leverage and Cook’s distance, see **Methods**; line in **Fig. 4A**). Adding fixed effects of biobanks to the model yielded only a marginal gain (incremental adjusted *R*^2^ = +1.7%). Further adding biobank-specific slopes improved penalized fit only slightly more (incremental adjusted *R*^2^ = +3.3%). These results suggest that trait skewness in the cohort, rather than other biobank-specific features, largely determines the estimated sign bias. This pattern held when we used different minor allele frequency thresholds when calculating sign bias, though the incremental benefit of biobank-specific factors is more pronounced at less stringent minor allele frequency cutoffs (**Fig. S6**; **Tables S10, S12**).

**Figure 4:**
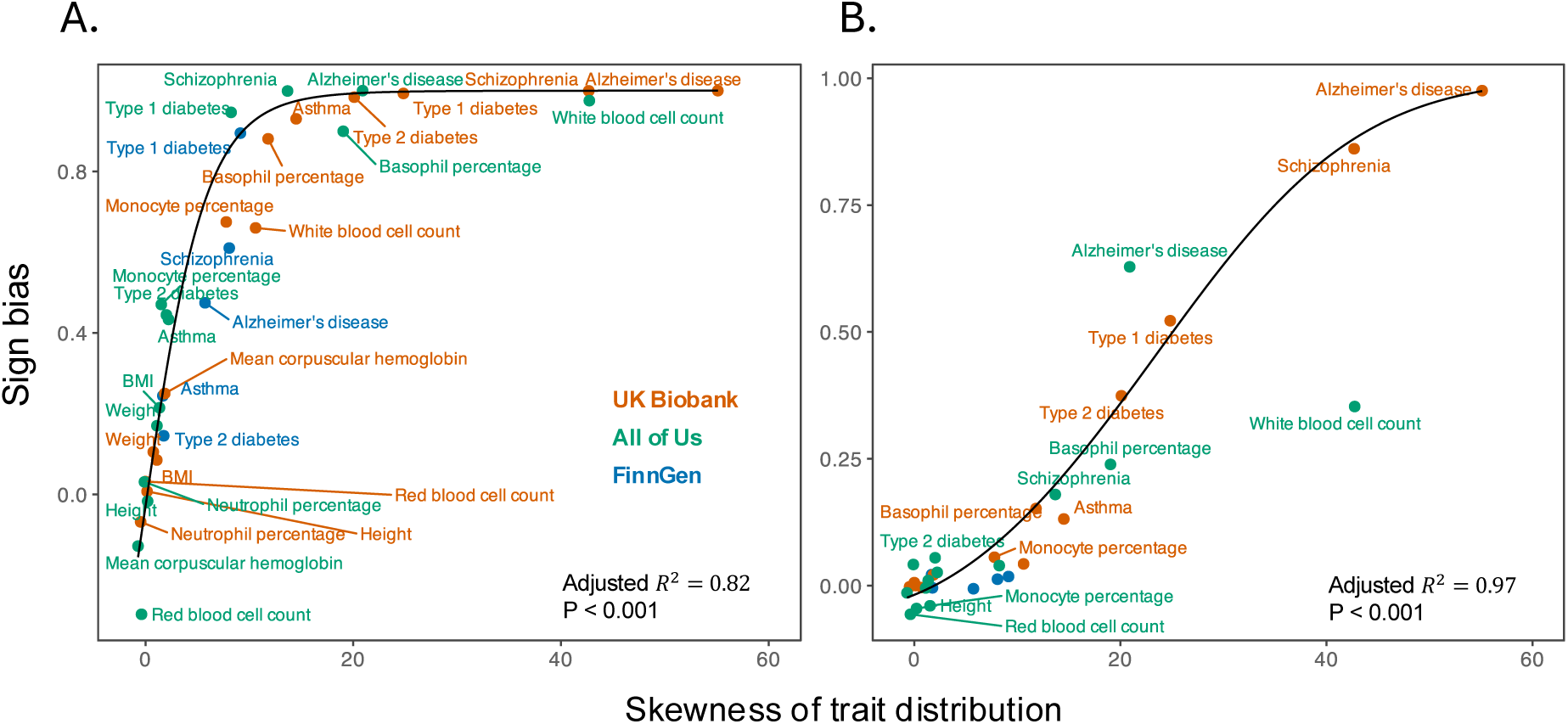
The skewness of trait distribution among study participants predicts sign bias estimates. Across traits and biobanks, the skewness (standardized third central moment) of the trait distribution predicts sign bias. Each point corresponds to a trait in one biobank. The sign bias here, as in Fig. 2B, was estimated across SNPs of minor allele frequency up to 0.1%. Morphometric traits clustered near zero skewness and zero sign bias, whereas imbalanced traits (Alzheimer’s disease, schizophrenia, type 1 diabetes) exhibited high skewness and near-unity sign bias. A pooled quadratic logit fit (black curve, with respective fit statistics shown in the lower right corner in each panel), ignoring biobank and trait identity, suggests that trait distribution skewness largely explains the relationships. White blood cell count is shown in the plot above but excluded from the regression fit because it is an influential high-leverage observation in AoU (see **Methods**). (A) using the most significantly trait-associated SNPs per approximately independent LD block; (B) using randomly selected SNPs in each approximately independent LD block.

We repeated the analysis using randomly sampled variants and again found a strong association between sign bias and trait skewness across traits and biobanks (pooled Spearman’s *ρ* = 0.843, *P <* 0.001; **Table S8**). The same quadratic logit model showed a tight fit (adjusted *R*^2^ = 0.97, *P <* 0.001 excluding the same outlier; **Methods**), and adding biobank-specific intercepts or slopes did not substantially improve the fit. Results were similar under alternative minor allele frequency cutoffs (**Fig. S7**; **Tables S10, S12**), with biobank-specific terms contributing more only at less stringent thresholds.

## Discussion

Our results suggest a link between the characteristics of genetic studies and inferences about genetic architecture. We studied sign bias as a case study of this link, and in this case large differences across biobanks were remarkably well explained by a simple summary of the distribution of the trait among study participants—namely, its skewness. This finding serves as a cautionary tale: summaries of genetic architecture can be influenced by characteristics of the study design, such as the recruitment mechanism, and do not necessarily reflect trait-level properties.

Differences in representation across studies are due in large part to differences in study design, such as how participants are recruited and enrolled into the study (e.g., population volunteers, patient-based enrollment, and disease-enriched designs^27–29)^, and how traits are measured. For example, participants recruited from clinical settings often have greater representation of participants with rare diseases, resulting in more cases in disease cohorts^65^.

For most traits, we observed lower SNP heritabilities in AoU compared to UKB. Chen et al.^16^ reported lower SNP heritabilities in other disease-enriched biobanks (e.g., BBJ) relative to population-based, volunteer cohorts such as UK Biobank and the Taiwan Biobank. To our knowledge, our results offer the first demonstration that the pattern of lower SNP heritability in disease-enriched biobanks holds even in ancestry-matched subsamples. Because power in GWAS depends directly on SNP heritability, one implication is that disease-enriched studies may be less powered to detect some genotype-trait associations compared with population-based cohorts.

What drives the difference in heritability? One possibility is biological differences between the cohorts, such as larger variance in effects not captured by common SNP variation (including environmental effects) among AoU participants compared with UKB participants. Another possibility is larger non-biological error in AoU, for example due to heterogeneity in trait definitions (e.g., disease classification and diagnostic codes) or measurement error. Many traits in AoU (and other disease-enriched studies) are derived from electronic health records and heterogeneous clinical laboratory data. These measurements are likely less standardized across study sites and time. This hypothesis is especially plausible for biomarkers and other clinical measurements, where differences in units or reporting conventions can add noise.

Although participation has been shown to affect estimates of genetic correlations^23^, the low correlations estimated for some traits (e.g., basophil percentage and type 1 diabetes) across biobanks are perhaps surprising, all the more so given our ancestry matching. These observations help contextualize comparisons of genetic architectures and polygenic score prediction accuracy^66,67^ across ancestry groupings. Some studies attribute discrepancies solely to differences in linkage disequilibrium (LD) patterns, causal allele frequencies or allelic turnover, or heterogeneity in the causal allelic effect distributions^68–70^. When cohort recruitment and other biobank characteristics are confounded with ancestry groupings, however, as they typically are, it becomes difficult to interpret differences in allelic effects (probed, e.g., via genetic correlation), heritabilities or the predictive utility of polygenic scores^71,72^. Furthermore, since recruitment and participation can differ by genetic ancestry groups within a biobank^73–75^, such factors may be confounded with ancestry in single-biobank analysis as well^71,76,77^.

Sign bias could also be affected by biological and evolutionary mechanisms beyond the effects of study design, such as biases in the sign of de novo mutations and natural selection. For example, quantitative traits can be skewed, even under neutrality, if mutational effects are asymmetric or heavy-tailed^78^. Additionally, a relationship between positive sign bias and allele frequency might be expected for traits (especially diseases) under purifying selection, such that risk-increasing mutations are constrained to lower frequencies^1,12,13,53,55,79^. This expectation matches our observations of mean sign bias across the minor allele frequency spectrum, which show stronger signals for disease traits. However, such mechanisms should be largely cohort-invariant and they do not explain why the same trait’s sign bias would differ across GWAS with similar sample composition of genetic ancestry. For an explanation based on natural selection to hold, selective pressures would have to differ dramatically across countries and produce detectable changes to genetic architecture over contemporary timescales and tiny population divergences^80–86^, which seems implausible. By contrast, our data show that cross-cohort differences in sign bias correlate with cohort-specific trait skewness (which is considerably higher for binary disease traits), rather than biobank identity per se. This relationship would be expected from a study design mechanism, in that properties of the trait distribution influence allelic effect estimation, thereby modulating the observed sign bias on top of any evolutionary baseline.

Throughout, we analyzed allelic effects using raw trait units of continuous traits rather than transforming them. In contrast, many GWAS analyses transform quantitative trait data (typically via inverse-rank normalization) so that the residuals are closer to conditionally Gaussian and measurement units are standardized across traits^87^. Such a transformation is useful for enhancing discovery power and producing “well-behaved” test statistics^88^. For example, inverse-rank normalization of the trait distribution would remove trait skewness and, as we show (**Figs. S8, S11**), result in little-to-no sign bias. It is therefore unsurprising that, to our knowledge, most studies have not found pronounced sign bias in allelic effects from GWAS for highly symmetric traits, transformed or not (see for example the symmetric “smile” plot of effect size vs. allele frequency for trait-increasing alleles reported by Koch et al^55^). However, transforming data (rather than using the scale deemed biologically relevant) can render allelic effect estimation and inferences about genetic architectures more mathematically convenient but less biologically interpretable^89,90^. In addition, we note that binary traits such as disease status are not typically transformed, and the interpretation of summaries such as sign bias on a linear liability scale may remain impacted by participation in genetic studies.

In this work, we challenge an often implicit assumption that inferences about the genetic architecture of traits and diseases are not biased by representation in genetic studies. We argue instead that such inferences are best viewed as cohort-dependent summaries rather than trait-intrinsic properties. Similarly, nominally “cross-ancestry” or “trans-ethnic” comparisons should be interpreted with caution, as apparent differences may partly reflect variation in recruitment or other aspects of study design. Going forward, it will be important to evaluate what facets of genetic architecture replicate across studies and why. It therefore seems vital to broaden the research community’s access to studies with diverse recruitment and participation profiles. This would help ensure that inferences about genotype-complex trait maps are generalizable and ultimately benefit everyone.

## Methods

### Data

We assembled a panel of traits that are present in both the UK Biobank (UKB) and All of Us (AoU) databases. The UKB cohort comprises approximately 500,000 residents of the United Kingdom aged 40-69 and recruited between 2006 and 2010^27^. The AoU program is a nationwide, longitudinal cohort in the United States aimed at recruiting participants from a diversity of ancestral backgrounds^28^. The database currently contains over 800,000 participants over 18 years of age, over 400,000 of whom currently have whole-genome sequences available. We chose to analyze 14 traits including three common morphometric traits (height, weight, and BMI), six hematological traits (monocyte percentage, basophil percentage, neutrophil percentage, white blood cell count, red blood cell count and mean corpuscular hemoglobin) and five diseases (asthma, type 1 diabetes, type 2 diabetes, schizophrenia, and Alzheimer’s disease).

For AoU, we used traits under the version 8 release in the Curated Data Repository (CDR). For physical measurements (height and weight), we queried the AoU CDR measurement table for records with “measurement source value” equal to “height” and “weight,” respectively. We restricted to rows with interpretable units by retaining measurements recorded in centimeters (height) and kilograms (weight) and excluded records with missing values or non-matching or unknown units. We then converted height from centimeters to meters and computed BMI per participant as weight (*kg*) divided by squared height (*m*^2^). For laboratory measurements, we queried the measurement table using the respective LOINC codes to match the corresponding units and measurement for the same trait in UKB (**Table S1**). For white blood cell percentages, we kept only records whose unit was a percentage (unit concept name in “percent”, “percentage unit”, “percent of white blood cells”). We excluded measurements with values lower than 0 or higher than 100. For white and red blood cell counts, we converted measurements to standard units. Specifically, white blood cell counts were standardized to thousands per microliter (10^3^*/µL*) and red blood cell counts were standardized to millions per microliter (10^6^*/µL*) using conversions for equivalent units (e.g., thousand per cubic millimeter, cells per microliter, and billion per liter). We excluded implausible values by removing negative measurements and applying specified bounds (white blood cell count: 0.1–500 in 10^3^*/µL*; red blood cell count: 0.5–12 in 10^6^*/µL*). For mean corpuscular hemoglobin, we retained only records with picogram-based units (unit concept name in “picogram”, “picogram per cell”, or “pg/cell”) and removed values outside bounds of 5–100 pg. With all laboratory traits, we kept only records with non-missing values and, because participants can have repeated measurements, we summarized each participant’s trait value as the median across all available measurements over time. For diseases in AoU, we pulled disease status by mapping ICD codes to phecodes using Phecode map X (https://phewascatalog.org/), requiring at least two distinct dates with a mapped code for case assignment.

To broaden the scope of the analysis, we also included matched disease endpoints from FinnGen, a nationwide Finnish research collaboration that connects genomic data (from genotype arrays and imputation) to decades of digital health registries in Finland for over 500,000 participants^29^. Since we did not have access to individual-level FinnGen data, we only used it in the analysis of sign bias presented in **Figs. 2** and **4** and further only used disease traits for which we can calculate the skewness based on sample prevalence. More details on the traits used are in **Table S1**.

### GWAS

For UKB, we downloaded the GWAS summary statistics from the Neale Lab (http://www.nealelab.is/uk-biobank/). These consisted of 13.7 million SNPs including both genotype array and imputed SNPs based on a subsample consisting of 361,194 unrelated individuals who cluster together in genetic principal component space (within a 7 standard deviation radius of individuals of “White British” ancestry) and who self-reported as White and British, Irish, or White ethnicity. For quantitative traits, we used summary statistics for the raw, untransformed traits. We additionally used the public release of FinnGen GWAS summary statistics (DF12 release), performed on 21,311,644 array and imputed SNPs using 500,348 samples for the 5 disease endpoints (https://r12.finngen.fi/). For all downstream analyses for UKB and FinnGen, we removed from the GWAS summary statistics all non-biallelic variants, variants with INFO scores *<* 0.8, variants with a Hardy-Weinberg exact test *P <* 1 × 10^−10^, and variants with a minor allele count ≤ 20 .

For AoU, we performed a GWAS on the same 14 traits, but sought to first identify a sample with similar ancestry composition to that used to generate the Neale Lab’s UKB-based summary statistics. We projected both the UKB and AoU individuals into the same PC space using pre-computed SNP loadings provided by the Global Biobank Meta-analysis Initiative^91^. We removed individuals who were flagged for QC issues or identified as close relatives (estimated kinship coefficient of more than 0.1, which covers first-and second-degree relatives) using summaries provided by AoU. We excluded individuals whose self-reported gender was not man or woman or whose self-reported sex was not male or female. We then trained various classifiers on the UKB cohort to predict White British-like ancestry using the first 10 PCs from the shared PC space. The random forest classifier (F1-score 0.97) was used to identify the ancestry-matched cohort from AoU (*n* = 221, 105). Self-reported race was used to validate that results indicate majority-similar self-reported race categories between the UKB and AoU samples (in this case ’White-British’ and ’White’), but were not used to determine inclusion in the matched AoU sample. Additionally, for each trait in our analysis, we removed individuals with missing values for that trait. We were left with between 68,492 to 199,529 AoU individuals for each trait.

We performed all GWAS using linear regression with PLINK 2.0^92^ on the raw trait values for each trait. We included the covariates age, age^2^, sex, sex×age, sex×age^2^, and the first 16 principal components of the genotype matrix of the ancestry-matched sample (all covariates were standardized to have mean 0 and variance 1 by using the PLINK flag --covar-variance-standardize). We used SNPs from the All of Us Allele Count/Allele Frequency (ACAF) subset of the whole-genome sequence data. We performed the analysis on a set of 10,102,149 matched autosomal variants (based on dbSNP rsIDs) in both the UKB and AoU datasets. We excluded variants with a call rate *<* 95%, and excess-heterozygosity outliers using a one-sided mid-p Hardy–Weinberg exact test (flags --midp and --keep-fewhet) with threshold *P <* 1 × 10^−12^. For all downstream analyses we only used summary statistics for biallelic SNPs and only kept variants with minor allele counts *>* 20 in the final variant set.

### Estimates of genetic architecture summaries

Using GWAS summary statistics, we estimated narrow-sense heritability using an additive SNP model for each trait and the genetic correlations between biobanks using LD Score Regression (LDSC)^33,47^. We calculated observed-scale estimates of SNP heritability for all traits, and further computed liability scale estimates for binary traits, assuming that the population prevalence matches the prevalence in the given biobank. We estimated effective polygenicity for each trait with S-LD4M^34^. For all analyses, we used pre-calculated LD scores for approximately 1.2 million HapMap3 variants and for the European ancestry group as provided by Bulik-Sullivan et al^33^.

### Estimating sign bias

For each trait, we polarized GWAS-estimated SNP allelic effects such that the sign represents that of the minor allele relative to the major allele. We estimated posterior sign probabilities for each SNP included in the GWAS using adaptive shrinkage (ash^58^). We estimated the posterior sign of the allelic effect, *η*, for variant *i* as

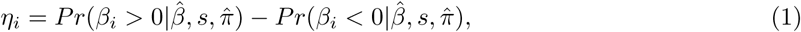

where *Pr*(*β_i_ >* 0|*β*^^^*, s, π*^) is the posterior probability (estimated using ash) that the minor allele of variant *i* is trait-increasing, and *Pr*(*β_i_ <* 0|*β*^^^*, s, π*^) is the posterior probability that it is trait-decreasing. In both terms, *β_i_* denotes the (unknown) true effect for SNP *i*, *β*^^^ is the vector of observed minor-allele effects estimated from all GWAS SNPs, *s* denotes the estimated standard errors of *β*^^^, and *π*^ denotes the empirical Bayes mixture weights over a point mass at 0 and a set of zero-centered Normal distributions with distinct variances. *η_i_*approaches +1 (−1) with increasing certainty that the minor allele is trait-increasing (decreasing) and nears 0 either when there is high uncertainty about the sign or the posterior probability that there is no effect is high.

The estimation of the sign bias in a given set of SNPs can be confounded by linkage disequilibrium (LD) among SNPs in a set. To avoid this confounding, we only use one SNP per approximately independent LD block following the protocol used in Zhu et al^90^. For the UKB, we used the 1,703 blocks for the European subset (EUR “superpopulation”) of the 1000 Genomes Phase 3 dataset^82^ as inferred by Berisa and Pickrell^93^. For AoU, we used the updated map of 1,361 approximately independent LD blocks for the EUR individuals for human genome build hg38 from MacDonald et al^94^.

We used two approaches to estimate the sign bias signal across loci: 1) across the most significantly trait-associated SNPs and 2) across randomly sampled SNPs. Because the magnitude |*η_i_*| quantifies certainty about the direction of the effect for variant *i* (|*η_i_*| = 1 indicates complete certainty; |*η_i_*| = 0 indicates maximal uncertainty), we can summarize the signal in a way that gives lesser weight to uncertain signs. For a set of SNPs *k*, we select a single variant (either at random or with the smallest association *p*-value) from each of *b* LD blocks, denote the resulting representative SNPs as 1*_k_, . . . , b_k_*, and average their sign estimates with weights proportional to their certainty as

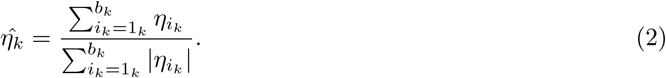

*η*^*_k_* approaches +1 when all the minor alleles in set *k* are trait-increasing, -1 when they are all trait-decreasing, and 0 when evidence is equal for trait-increasing and trait-decreasing effects. Within each set *k*, we excluded LD blocks that contained fewer than 10 SNPs. In the main text analysis, we define a SNP set *k* either as all variants within a range of minor allele frequencies or all variants below a specified allele frequency threshold.

In the significance-based approach, we calculated standard errors via bootstrapping. For each variant set *k*, we resampled the *b* contributing LD blocks with replacement, selected the SNP with the lowest *p*-value within each resampled block, and recomputed **Eq. 2**. We repeated this procedure 1,000 times and used the standard deviation of these values as the standard error. In the randomized approach, we quantified uncertainty by repeated random selection. For each set *k*, we randomly selected one SNP within each contributing LD block, computed **Eq. 2**, and repeated this procedure 1,000 times. We used the mean of these 1,000 values as the point estimate, and the standard deviation across replicates as its standard error.

### Simulating traits and associated allelic effects

To understand how trait skewness can drive sign bias, we carried out two simulation schemes. In both schemes, we simulated genotypes and traits in a population, performed GWAS in samples chosen with varying trait skewness, and examined downstream GWAS summary statistics. This allowed us to isolate mechanisms by which trait distribution skewness can generate inferred sign bias despite there being no sign bias in the generative mutational process.

The first scheme (scheme A) produces increasingly positive cohort trait means and associated trait skewness by sampling from increasingly higher trait values within a unimodal population distribution. We simulated a population of *N* = 2 × 10^6^ individuals with equal numbers of independent trait-increasing and trait-decreasing alleles of equal magnitude, consisting of *M* = 4,000 total variants. For each of *M/*2 matched pairs, we assigned each variant *j* an allele frequency *p_j_* log-uniformly over a specified range from *p_j_* = 1 × 10^−4^ to *p_j_* = 0.1 and assigned it to both variants. We assigned each variant *j* a true allelic effect 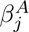 with common magnitude 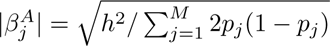 , with *h*^2^ = 0.6. This procedure ensured that the effect magnitude was constant across variants and independent of allele frequency, leading to no sign bias in the population. We drew genotypes for each individual under Hardy-Weinberg equilibrium to yield genotype dosages 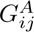 as

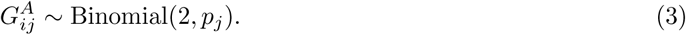

We assumed independence among variants (no linkage disequilibrium) and individuals (no kinship). Adding independent environmental noise 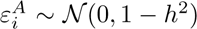 , we assigned individual *i*’s trait value, 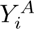 as

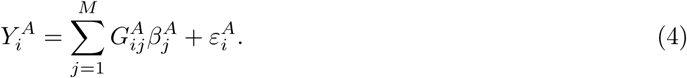

We aimed to create cohorts in which sampling is related to the trait value such that the cohort’s trait distribution can differ from the population distribution in a controlled way. Specifically, we wished to vary how strongly the cohort is enriched for higher (or more positive) trait values. To achieve this goal, we drew cohorts from this population as follows. We first divided the population of *N* individuals into *V* = 200 quantile bins based on their trait values such that each bin contains *N/V* = 10, 000 individuals. The quantile binning method allows us to sample these bins on the rank scale rather than the raw trait scale and compare across cohorts whose trait distributions differ. We use the midpoint of a quantile bin (rather than its endpoints) to represent the bin’s location on the [0, 1] rank scale. For a bin *v* ∈ 1*, . . . , V* , we define its mid-quantile rank as

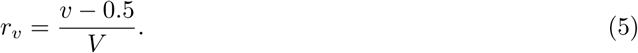

For each cohort, we imposed different cutoff values *q* ∈ [0, 1) so that a given cohort is sampled from bins with *r_v_* ≥ *q*. Increasing *q* restricts sampling toward increasingly larger values of the trait. We apply bin weights to simulate sampling within the retained portion of the distribution. Among these retained bins *V_q_*, we assigned each bin a weight

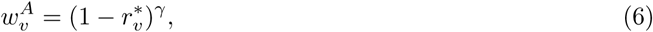

where

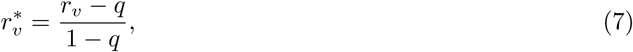

and *γ* is a shape parameter controlling how strongly the sampling concentrates near the cutoff *q* (larger *γ* places more mass on bins just above *q*). To avoid drawing an entire cohort from a narrow set of bins, we used a mixture sampling rule: we drew a fraction *τ* of a cohort uniformly across the retained bins, and we drew the remaining fraction 1 − *τ* according to the weighted distribution proportional to 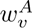 . Equivalently, the probability that a randomly selected cohort member is assigned to bin *v* ∈ *V_q_* is

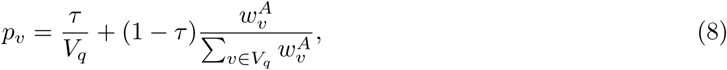

and *p_v_* = 0 for *v* ∈*/ V_q_*. Given *p_v_* and a cohort size of *n* individuals, we drew bin sample counts as

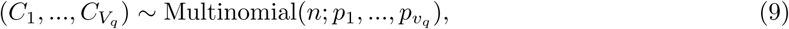

so that *C_v_* is the number of sampled individuals assigned to bin *v*. We then sampled exactly *C_v_* individuals without replacement from the set of individuals in bin *v*, and concatenated sampled individuals across bins to form a cohort.

We calculated cohort trait skewness as the standardized central third moment of its distribution and computed both the “true” sign bias in a cohort (from the known, simulated causal effects) and estimated sign bias (from GWAS summary statistics) for all variants with sample minor allele frequencies up to 1%. To calculate the true sign bias, we oriented each SNP’s true effect to the minor allele and averaged across the true sign of effects. To estimate sign bias, we performed genome-wide association analyses (GWAS) within a cohort on each variant using ordinary least squares regression. As described in the section **Estimating sign bias**, we performed adaptive shrinkage (ash) on all variants and estimated per-variant sign bias using **Eq. 1**. Because our main analyses focus on subsets of variants chosen for statistical significance, we partitioned SNPs (in order of their assigned index) into consecutive groups of six and selected the single SNP with the smallest two-sided GWAS *p*-value in each group. We then used this set of selected variants to compute the aggregate certainty-weighted cohort sign bias following **Eq. 2**.

Using the above approach, we generated cohorts of *n* = 10, 000 sampled from a single simulated population spanning a range of skewness values (target skewness values ranging from 1 to 10), creating 20 replicate cohorts for each target skewness value. We held (*γ, τ* ) fixed at *γ* = 20 and *τ* = 0.01 across all replicates and tuned only the cutoff *q* via a grid search from 0 to 0.95 to identify values producing cohort skewness closest to each target.

In the second scenario (scheme B), we added latent subgroup structure to test whether the same sampling logic produces qualitatively similar results when the population trait distribution is multi-modal. Namely, we constructed cohorts and associated skewness from a tri-modal population by sampling individuals near the centers of the central and right modes and varying the proportion of individuals from the right mode sampled into the cohorts. We made the population distribution symmetric overall so that any directional shift in the cohort trait distribution arises from the sampling process. Here, we generated the population by assigning each individual *i* to a latent mode *Z_i_* ∈ *L, C, R* (left, center, and right) with

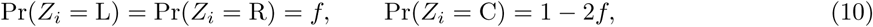

where *f* is a fixed tail fraction equal to 0.1. This fraction keeps most individuals in the center mode but ensures that a substantial proportion of individuals are present in the left and right modes.

We simulated paired variants as in scheme A using the same number of loci (*M* = 4, 000) and equal magnitudes of effects, with *h*^2^ = 0.6. We assigned initial allele frequencies in each variant pair as in scheme A. However, to separate the latent modes, we adjusted these allele frequencies to depend on both mode and effect sign. We included a shift parameter *δ* to make the left, center, and right modes genetically different by increasing trait-increasing allele frequencies in the right mode and trait-decreasing allele frequencies in the left mode. If the effect size of variant *j* is 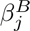 , let 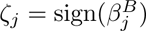 and define *t*(L) = −1, *t*(C) = 0, and *t*(R) = +1. We set the mode-specific allele frequency for variant *j* as

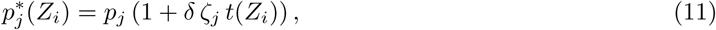

with shift parameter set to *δ* = 0.5. We chose this value to induce substantial allele frequency differentiation between modes while keeping frequencies within a valid range given our constraint of *p_j_*≤ 0.1.

Genotypes were then drawn conditional on mode as

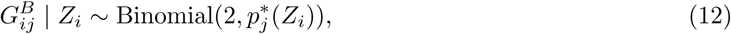

and a trait *Y ^B^* was generated as in scheme A (see **Eq. 4**).

Cohorts of size *n* = 10, 000 were then sampled using a two-component rule: a fraction *ρ* was drawn from the right mode (*n_R_*= *ρn*) and the remainder from the center mode (*n_C_* = *n* − *n_R_*). Varying *ρ* controls the mixture composition of the resulting cohort and therefore provides a direct way to tune the skewness of the cohort. Within each mode *z* ∈ {*C, R*}, sampling was restricted to a narrow region around the mode-specific median trait value. Let *m_z_* be the median of trait *Y ^B^*|*Z* = *z* and *s_z_* be the standard deviation of *Y ^B^*|*Z* = *z* so that for individuals with *Z_i_*= *z*

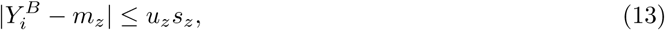

where *u_z_* defines how wide the sampling window is around mode *z*’s median. In the present simulation scheme, *u_C_*= 0.1 and *u_R_*= 0.037. These values create narrow windows that concentrate sampling near each mode’s center while retaining enough eligible individuals. If we let *λ_z_* be a parameter that controls how strongly sampling within mode *z* concentrates near the median, individuals inside each of these sampling windows are then given weights

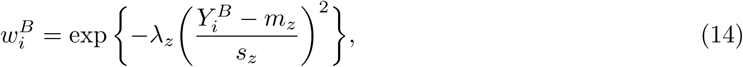

with *λ_C_* = 200 and *λ_R_* = 2. We used a large *λ_C_* to approximate highly concentrated sampling around the median of the center mode, and a smaller *λ_R_* to allow greater within-mode variation in the right mode. Within each mode’s sampling window, we then sampled individuals without replacement with probability proportional to 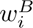 to create cohorts. We varied only the right-mode fraction *ρ* to reach target skewness values of 1–10 (all other sampler parameters were fixed), and generated 20 replicate cohorts for each target skewness value (**Table S6**). All remaining steps for computing skewness as well as true and estimated sign bias were performed as in scheme A.

In schemes A and B above, populations contain an equal number of causal trait-increasing and trait-decreasing variants, each with the same absolute effect size. In a third simulation, we investigated whether cohort trait skewness could produce a similar pattern in the absence of causal effects. To do so, we repeated all the steps outlined above for simulation scheme A, but with all variant effects set to zero. We then estimated the sign bias from the resulting GWAS summary statistics using the same procedures described above (**Fig. S11**).

### Evaluating the relationship between trait skewness and sign bias

We processed traits in the UKB to correspond to their GWAS equivalents by following the sample QC procedures outlined by the Neale Lab (for the UKB v3 GWAS; http://www.nealelab.is/uk-biobank/). For AoU, we used the trait distribution for individuals who passed our sample QC steps outlined above in **GWAS**. For FinnGen disease endpoints, information about the number of cases and non-cases used in each GWAS is publicly available^29^. For each trait distribution, we calculated the skewness as the standardized third moment. We did not apply any normalization or covariate residualization; skewness is evaluated on the observed scale within each biobank.

We paired the computed skewness of the raw traits with sign bias estimates for alleles with frequencies ≤ 0.1% (these estimates are shown in **Fig. 2B** and **Fig. S4** calculated from **Eq. 2**). We denote the sign bias estimate for a given trait *T* in biobank *d* as *η*^*_T,d_*. Because our sign bias estimates are naturally bounded between -1 and 1 and represent differences of posterior sign probabilities, we map *η*^*_T,d_* to a probability scale *p*^*_T,d_* = (*η*^*_T,d_* + 1)*/*2, which corresponds to the posterior probability that the minor allele effect is positive. We place *p*^*_T,d_* on an unbounded scale using the logit *ℓ_T,d_* = logit(*p*^*_T,d_*), which improves the stability of the variance for probabilities close to 1.

We then modeled the association between trait skewness and sign bias with a second-degree polynomial

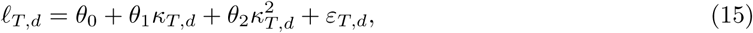

where *κ_T,d_* denotes the standardized third central moment (skewness) of trait *T* in biobank *d*, *θ*_0_ is the global intercept, *θ*_1_ and *θ*_2_ are the global slope terms, and *ε_T,d_* are mean-zero residuals.

We additionally fit a model that allowed for biobank-specific intercept shifts:

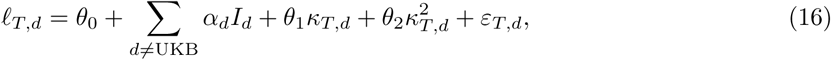

where *I_d_* acts as an indicator for biobank *d* (AoU or FinnGen, with UKB used as a reference), and *α_d_* is the shift in intercept for biobank *d*.

Finally, we fit a model that allowed for biobank-specific interactions:

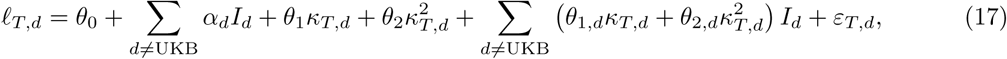

where *θ*_1*,d*_ and *θ*_2*,d*_ capture biobank-specific deviations from the UKB slope terms.

We computed standard influence diagnostics for each observation (*T, d*) under the interaction model and removed observations from the primary fits that were simultaneously very high leverage and highly influential on the regression coefficients. To do so, we applied thresholds for both leverage and Cook’s distance to define the exclusion criteria^95–97^. Specifically, we excluded observations with leverage values both exceeding 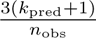 and Cook’s distance values twice the conventional threshold of 4*/n*_obs_. Here, *k*_pred_ is the number of predictors in the regression (excluding the intercept) and *n*_obs_ is the number of observations used to fit the model. This criterion removed only white blood cell count in AoU (leverage ≈ 0.99; Cook’s distance ≈ 0.34). Including white blood cell count lowers model fit (adjusted *R*^2^ = 0.76, *P <* 0.001 for alleles up to frequencies of 0.1%) and substantially increases the incremental benefit of adding biobank-specific slopes (incremental adjusted *R*^2^ = +10.2%), indicating that differences between biobanks are sensitive to this single trait (**Fig. S6**; **Tables S11, S12**). This sensitivity is biologically plausible as white blood cell count can change substantially depending on transient factors (e.g., acute infection/inflammation, stress responses, or medication effects)^98^. Whereas UKB used more standardized protocols across assessment centers to assess total blood counts, this trait in AoU is largely EHR-ascertained across multiple health care organizations with heterogeneous collection contexts. Such a collection approach is likely to combine diverse physiological states, including baseline values with episodic inflammatory “spikes,” potentially contributing to its varying skewness across cohorts.

## Acknowledgments

We thank the National Institutes of Health All of Us Research Program, the UK Biobank, FinnGen, and their participants for their essential contributions and for providing access to the participant data used in this research. This study used data from the All of Us Research Program’s Controlled Tier Dataset v8, available to authorized users on the Researcher Workbench. We thank Doc Edge, Lin Poyraz, Shaila Musharoff, Mark Kirkpatrick, Molly Przeworski, Nasa Sinnott-Armstrong, Guy Sella, Jeff Spence, Elliot Tucker-Drob, Yuval Simons and members of the Harpak Lab for helpful discussions and/or comments on the manuscript. This work was supported by NIH grant R35GM151108 and a Pew Biomedical Scholarship to A.H. This research used the UK Biobank resource (application 92741) and was approved by The University of Texas at Austin Institutional Review Board (Protocol 00003287).

## Data and code availability

Data and code generated by this study can be found at 10.5281/zenodo.1822803799. The code used to generate all results and figures is also deposited at https://github.com/harpak-lab/sign_bias_gwas.

## Notes

http://www.nealelab.is/uk-biobank/

https://r12.finngen.fi/

### Competing Interest Statement

A.H. is a paid strategic advisor for Yuimedi Corporation.

### Summary of Updates

Additional simulation analysis incorporated; Figure 1 revised for clarity; Supplemental files updated; and edits made to Results and Discussion sections in the text.

