## Supplemental Text and Figures for "Representation in genetic studies affects inference about genetic architecture"

### Supplementary text

#### Text S1: Comparison of genetic correlations across datasets

To verify that departures from a genetic correlation of 1 across AoU and the Neale UKB summary statistics (from `http://www.nealelab.is/uk-biobank/`) are not driven primarily by sample estimation error, we constructed a within-cohort benchmark by splitting the UKB into non-overlapping subsets, repeating GWAS in each subset, and estimating genetic correlations between subsets using the same `LDSC`[1] pipeline.

We used the same UKB phenotypes described in the main text (under Data in **Methods** in main text). We pulled individual level data for height (field 50), weight (field 21001), BMI (field 21002), monocyte percentage (field 30190), neutrophil percentage (field 30200), basophil percentage (field 30220), white blood cell (leukocyte) count (field 30000), red blood cell (erythrocyte) count (field 30010), and mean corpuscular hemoglobin (field 30050). For binary traits, we used individual level data from field 41202 using three-character ICD10 roots: Alzheimer's disease (G30), Asthma (J45), type 1 diabetes (E10*), type 2 diabetes (E11*), and schizophrenia (F20*). We curated the samples using the provided QC fields and genotype-derived principal components, following the steps outlined by the Neale lab, excluding individuals flagged as heterozygosity/missingness outliers (field 22027), showing putative sex chromosome aneuploidy (field 22019), that were excluded from kinship inference or with excess relatives (field 22021), and were not used in the computation of UKB principal components (PCs; field 22020). We used the curated White British subset (22006=1) passing base QC to define the reference PC distribution. Using the first six genetic PCs (field 22009; renamed PC1-PC6), we computed each individual's squared standardized distance from the White British centroid and retained individuals within a 7 SD ellipse. We further filtered for self-reported "White" ethnicity (field 21000 with values 1, 1001, 1002, or 1003). The final GWAS sample contained 360,951 individuals. We then split this sample randomly into two groups, one with 180,475 individuals (UKB1) and one with 180,476 individuals (UKB2). Details of the resulting traits used in the GWAS are given in **Table S13**.

For each sample, we ran GWAS on the 14 traits using `PLINK` 2.0[2] (linear regression for quantitative and binary traits using `--glm`) on the imputed autosomal variants. We excluded variants with a minor allele frequency below 0.1%, a missing genotype rate > 5%, and with a Hardy-Weinberg exact test $P < 1\times10^{-10}$ (using `--midp` and `--keep-fewhet flags`). All models included an intercept and adjusted for standard covariates, including age, age$^2$, sex, sex×age, and PCs 1-20. All covariates were standardized to mean 0 and variance 1 (using the flag `--covar-variance-standardize`). We were left with 15,653,401 variants in the first group (UKB1) and 15,653,708 in the second (UKB2).

We used bivariate LD score regression[1] to compare the genetic correlations between groups UKB1 and UKB2 across traits, and also compared each UKB group with our summary statistics from AoU (**Fig. S9**). We used precalculated LD scores of HapMap3 variants for the European ancestry group for all analyses[1].

### Supplementary figures

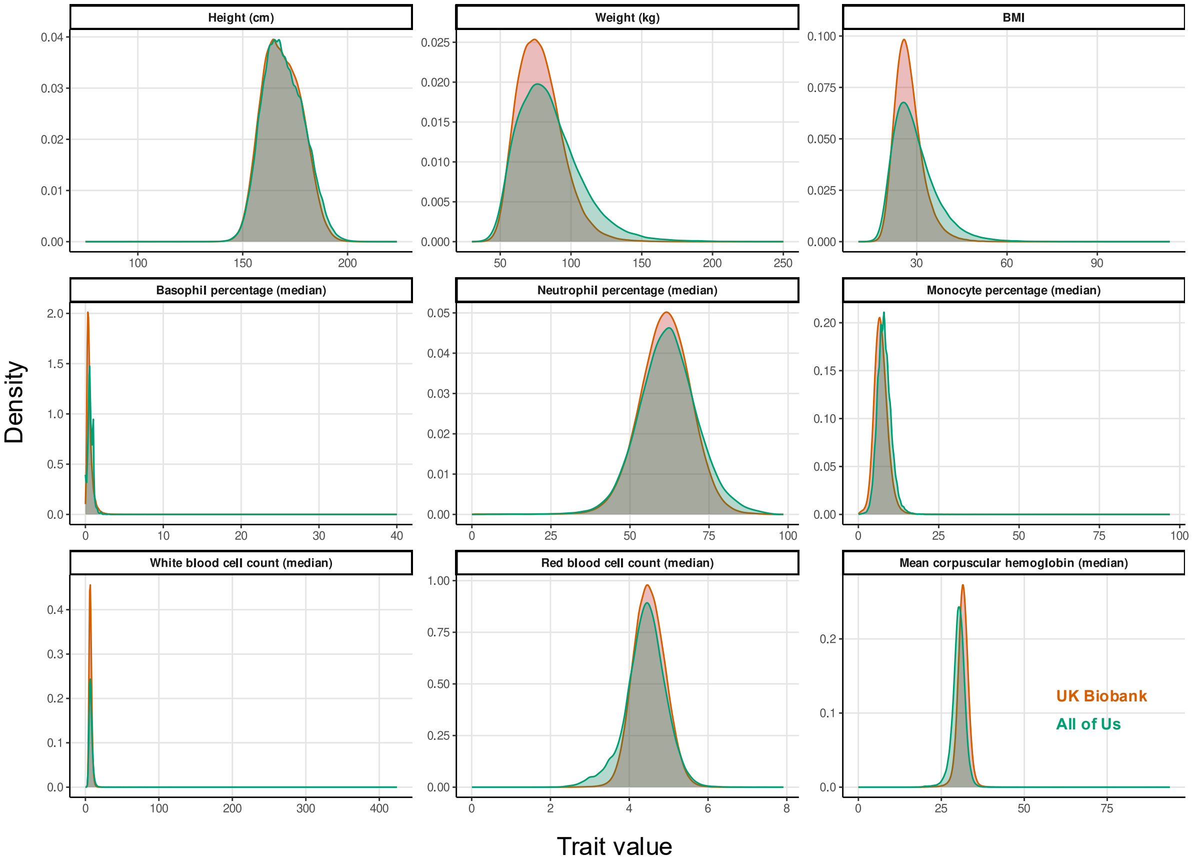

**Figure S1: Quantitative phenotype distributions in UKB and AoU.** The Y axis indicates the density of the distribution, and the X axis indicates the trait value. The orange density curves are the distributions in UKB and the green density curves are the distributions in AoU.

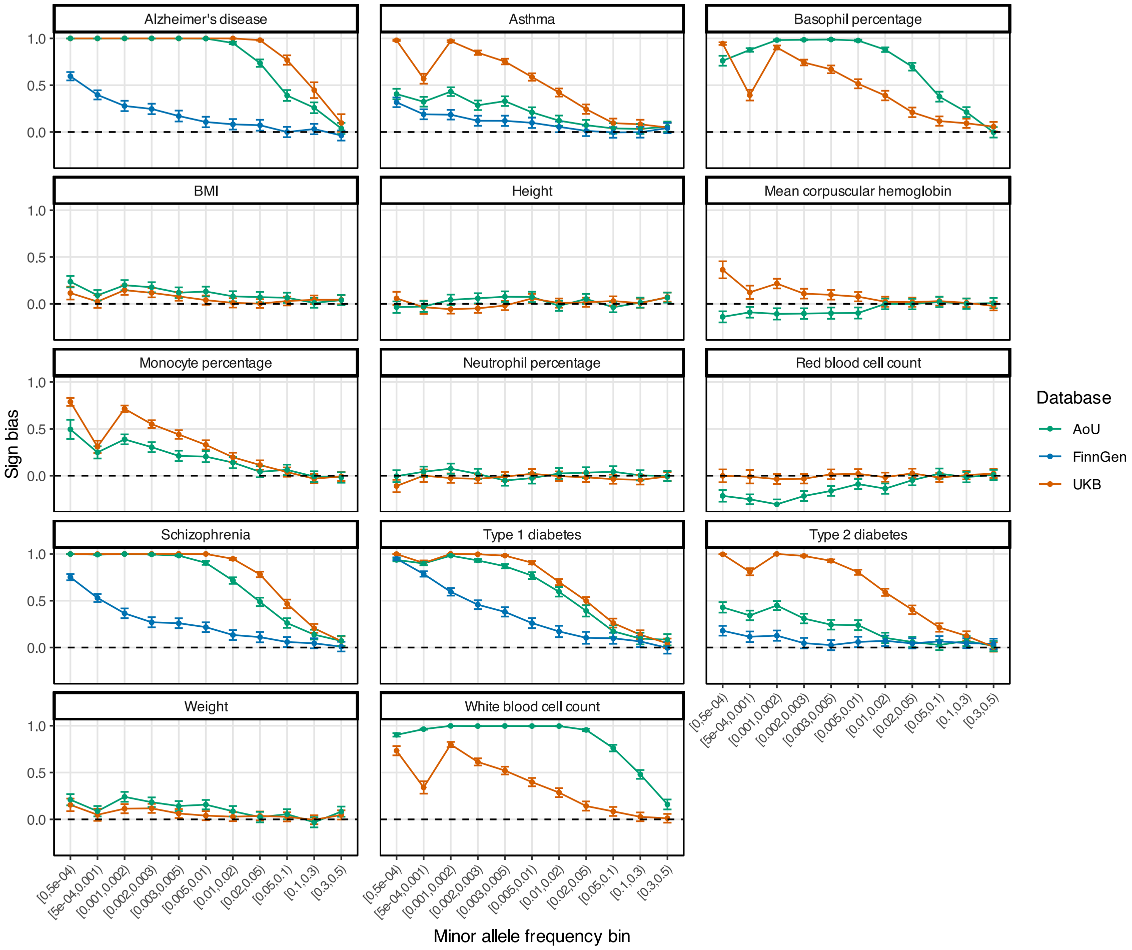

**Figure S2: Sign bias within each minor allele frequency bin for the most significant SNPs.** This figure mirrors the analysis of **Fig. 2A**, except that all analyzed traits are shown. For each trait and a range of minor allele frequencies, we considered the sign of one SNP per approximately independent LD block. Here, we specifically used SNPs that were the most significantly associated with the trait in the block. The sign bias for each set of SNPs ranges from -1 (all SNPs are trait/risk decreasing) to 1 (all SNPs are trait/risk increasing). The sign bias is estimated using empirical Bayes adaptive shrinkage (`ash`) to account for measurement uncertainty. Points show estimates and error bars show the 95% confidence intervals. Dashed horizontal lines show no sign bias (i.e., a value of 0).

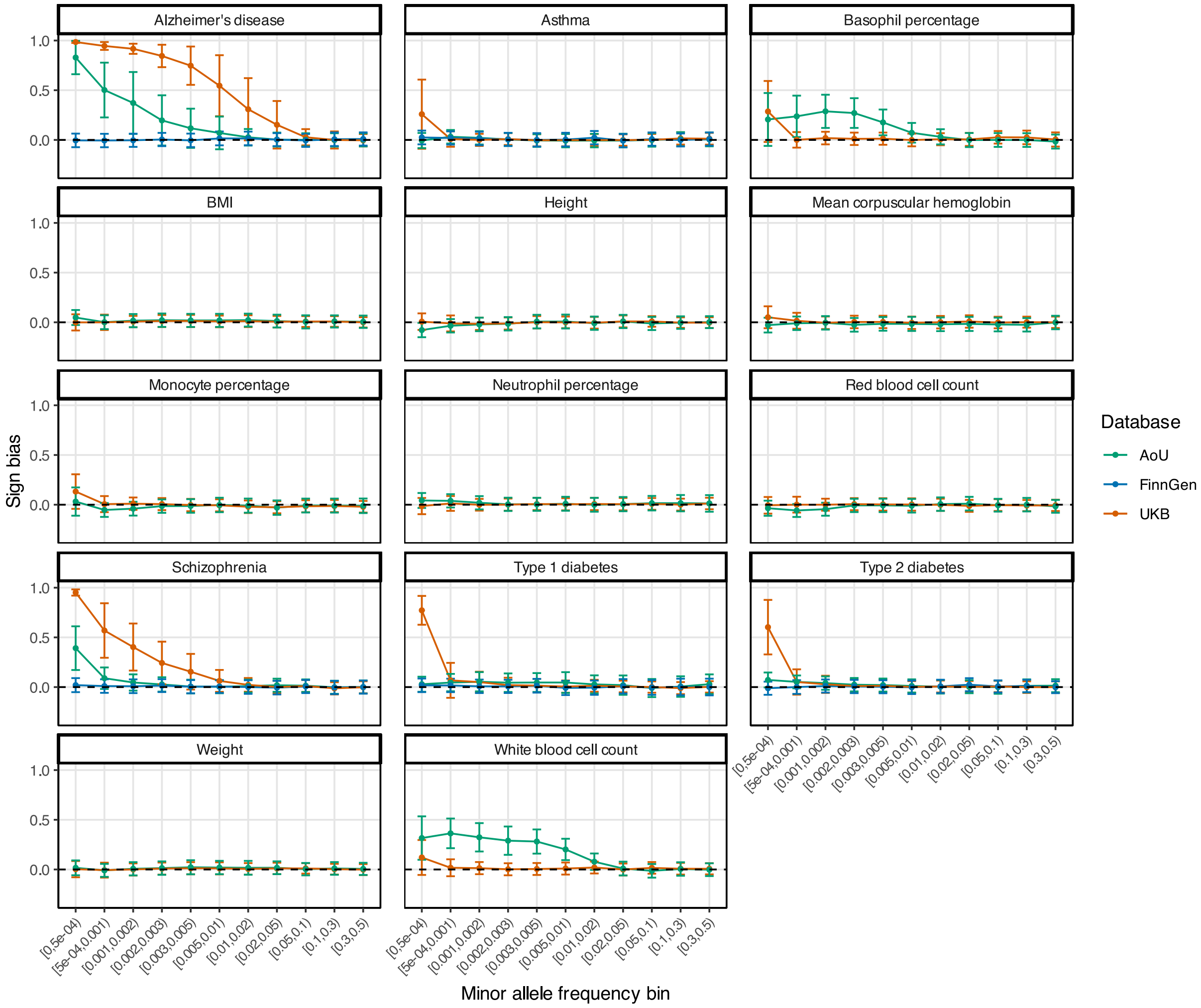

**Figure S3: Sign bias within each minor allele frequency bin for randomly sampled SNPs.** This figure mirrors the analysis of **Fig. 2A**, except that a variant was sampled at random from each LD block. This was done 1,000 times per bin and the mean across replicates was used as the estimate for each bin. Points show estimates and error bars show the 95% confidence intervals. Dashed horizontal lines show no sign bias (i.e., a value of 0).

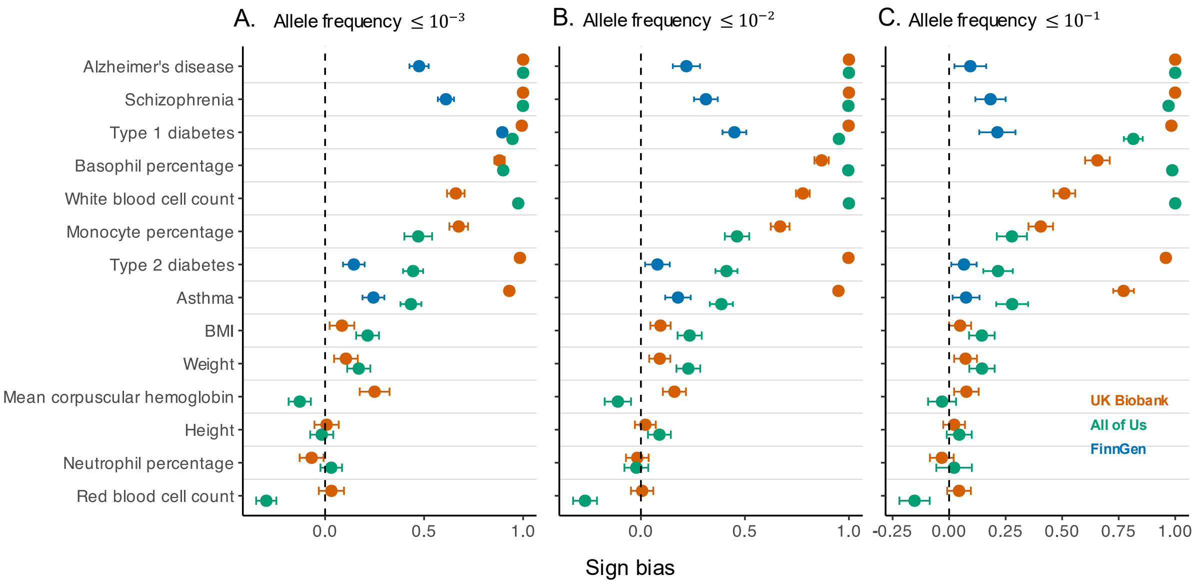

**Figure S4: Sign bias estimates across biobanks for the most significantly associated SNPs at different allele frequency thresholds**. This figure mirrors the analysis of **Fig. 2B**, except that different allele frequency thresholds are shown. The sign bias for each set of SNPs ranges from -1 (all SNPs are trait/risk decreasing) to 1 (all SNPs are trait/risk increasing). The sign bias is estimated using empirical Bayes adaptive shrinkage (`ash`) to account for measurement uncertainty. Points show estimates (UKB in orange, AoU in green, and FinnGen in blue) and error bars show the 95% confidence intervals. Dashed vertical lines show no sign bias (i.e., a value of 0). (A) Estimates for SNPs with minor allele frequency up to 0.1%, same as **Fig. 2B**. (B) Estimates for SNPs with minor allele frequency up to 1%. (C) Estimates for SNPs with minor allele frequency up to 10%.

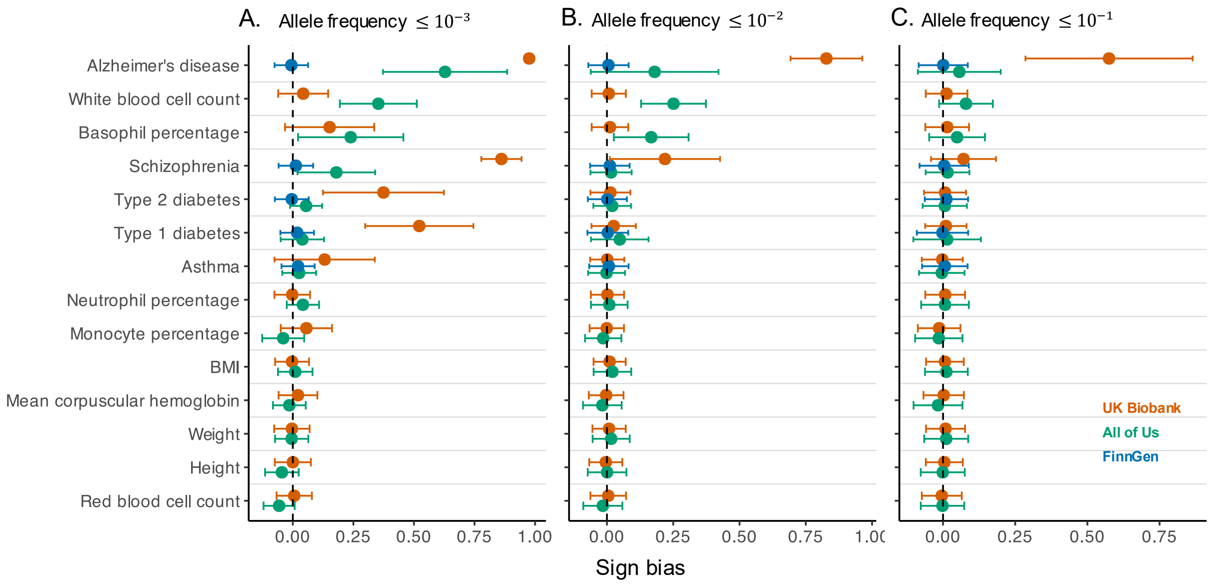

**Figure S5: Sign bias estimates across biobanks for randomly sampled SNPs at different allele frequency thresholds.** This figure mirrors the analysis of **Fig. S4**, except that a variant was sampled at random from each LD block. This was done 1,000 times per threshold and the mean across replicates was used as the estimate. The sign bias for each set of SNPs ranges from -1 (all SNPs are trait/risk decreasing) to 1 (all SNPs are trait/risk increasing). The sign bias is estimated using empirical Bayes adaptive shrinkage (`ash`) to account for measurement uncertainty. Points show estimates (UKB in orange, AoU in green, and FinnGen in blue) and error bars show the 95% confidence intervals. Dashed vertical lines show no sign bias (i.e., a value of 0). (A) Estimates for SNPs with minor allele frequency up to 0.1%, using the frequency threshold used in **Fig. 2B**. (B) Estimates for SNPs with minor allele frequency up to 1%. (C) Estimates for SNPs with minor allele frequency up to 10%.

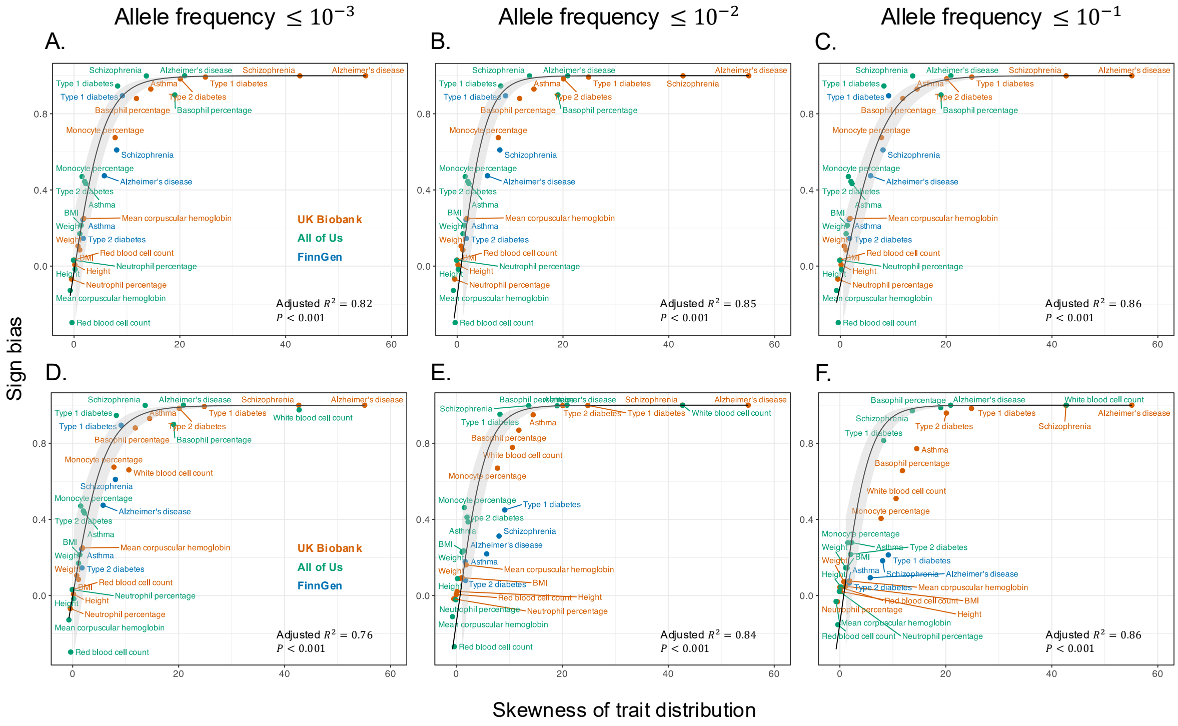

**Figure S6: The skewness of trait distribution among study participants predicts sign bias estimates using the most significantly associated SNPs.** This figure mirrors the analysis of **Fig. 4A**, except that different minor allele frequency (MAF) thresholds are shown. The skewness (standardized third moment) of the trait distribution predicts the sign bias, selecting the most significantly trait-associated SNP per LD block. Each point is a trait in one biobank (UKB in orange, AoU in green, and FinnGen in blue). The black curve (gray 95% CI) is a pooled quadratic-logit fit. *Top row (A–C)*: White blood cell count is excluded from the plots and the regression fits because it was flagged as a highly influential and high-leverage observation. (A) MAF $\leq 0.1\%$ (adjusted $R^2 = 0.82$, $P < 0.001$). (B) MAF $\leq 1\%$ (adjusted $R^2 = 0.85$, $P < 0.001$). (C) MAF $\leq 10\%$ (adjusted $R^2 = 0.86$, $P < 0.001$). *Bottom row (D–F)*: MAF thresholds correspond to (A–C), but white blood cell count is included in both the plots and the regression fits. (D) Adjusted $R^2 = 0.76$, $P < 0.001$. (E) Adjusted $R^2 = 0.84$, $P < 0.001$. (F) Adjusted $R^2 = 0.86$, $P < 0.001$.

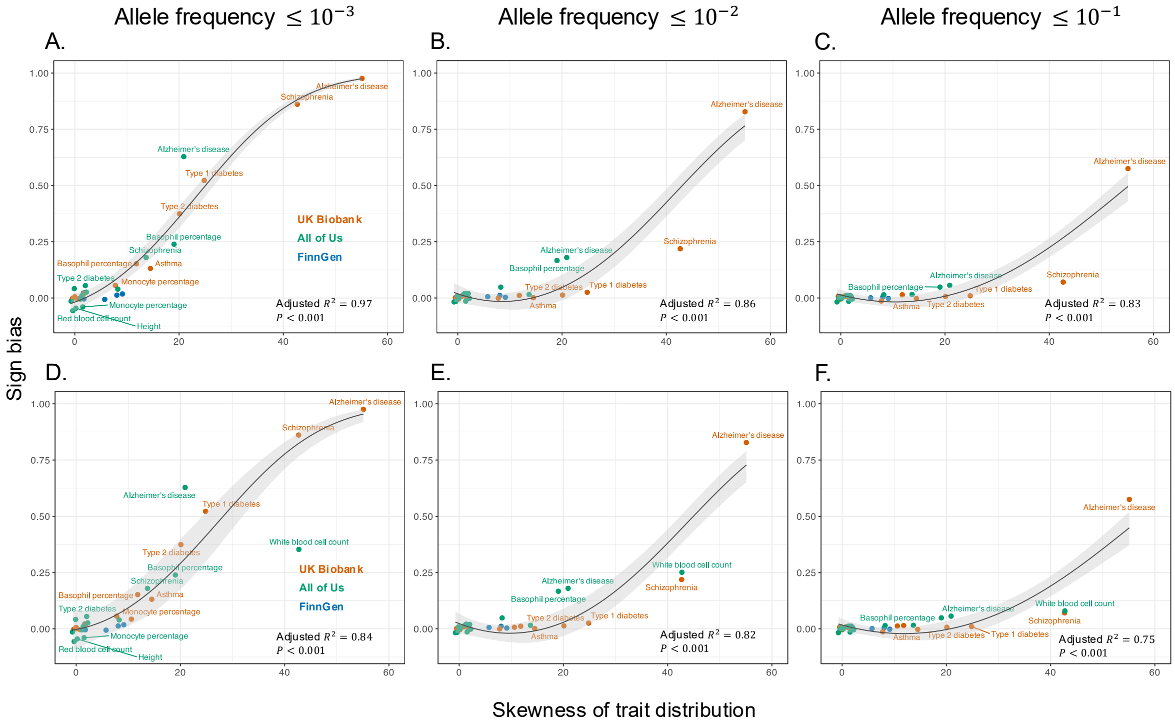

**Figure S7: The skewness of trait distribution among study participants predicts sign bias estimates using random SNPs.** This figure mirrors the analysis of **Fig. 4B**, except that different minor allele frequency (MAF) thresholds are shown. Each point is a trait in one biobank (UKB in orange, AoU in green, and FinnGen in blue). The black curve (gray 95% CI) is a pooled quadratic-logit fit. *Top row (A–C)*: White blood cell count is excluded from the plots and the regression fits because it was flagged as a highly influential and high leverage observation. (A) MAF $\leq 0.1\%$ (adjusted $R^2 = 0.97$, $P < 0.001$). (B) MAF $\leq 1\%$ (adjusted $R^2 = 0.86$, $P < 0.001$). (C) MAF $\leq 10\%$ (adjusted $R^2 = 0.83$, $P < 0.001$). *Bottom row (D–F)*: MAF thresholds correspond to (A–C), but white blood cell count is included in both the plots and the regression fits. (D) Adjusted $R^2 = 0.84$, $P < 0.001$. (E) Adjusted $R^2 = 0.82$, $P < 0.001$. (F) Adjusted $R^2 = 0.75$, $P < 0.001$.

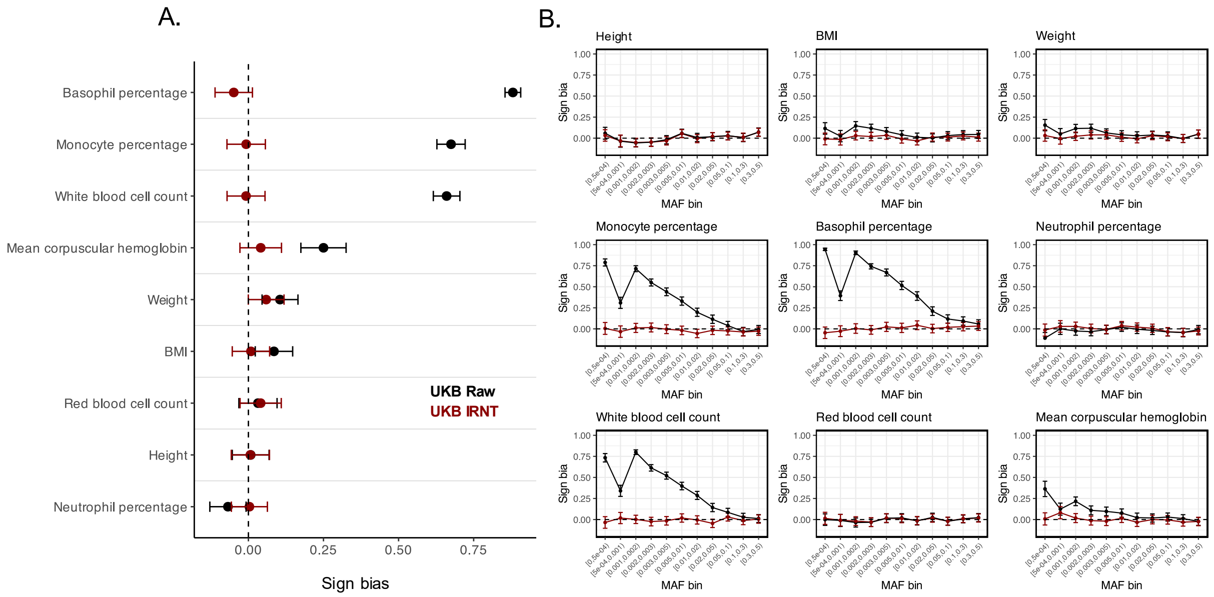

**Figure S8: Effect of inverse rank-normal transformation (IRNT) on sign bias estimates in UK Biobank.** We repeated the sign bias analysis described in the **Methods** and shown in **Fig. 2** in UK Biobank using GWAS performed on the IRNT-transformed quantitative traits (dark red) and compared them to results on the raw, untransformed traits (black). As in **Fig. 2**, we performed the analysis by selecting the most significantly trait-associated SNP per LD block. (A) Sign-bias estimates restricted to rare minor alleles with frequencies $\leq 0.001$, shown for each quantitative trait. (B) Sign-bias estimates stratified by minor-allele-frequency bins (x-axis) for the same traits. Points/lines show the estimated sign bias in each setting (with uncertainty shown by error bars; see **Methods**). Across traits, especially those with strongly skewed raw phenotypes, IRNT largely eliminates or attenuates the apparent sign bias relative to the raw-scale analysis. Because IRNT changes the trait scale, this attenuation cannot on its own distinguish genuine sign bias in genetic effects from that induced by biased estimation.

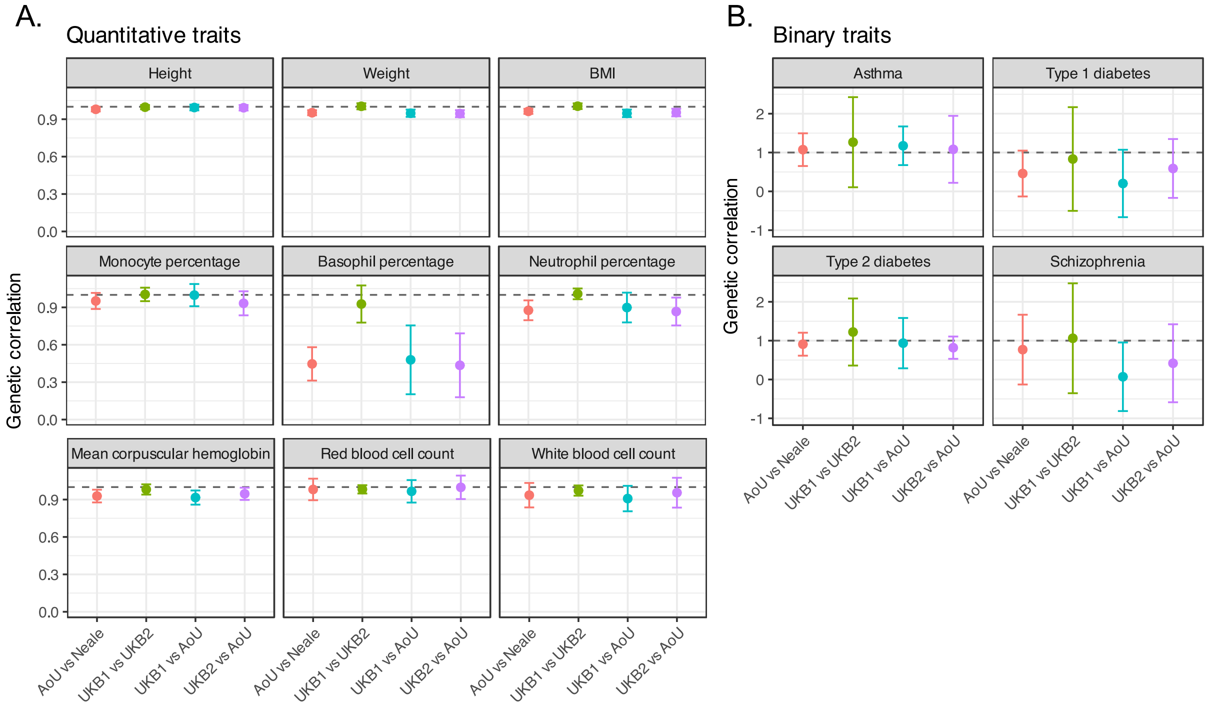

**Figure S9: Testing the expectation of perfect genetic correlations under the null.** Our analysis in **Fig. 1C** implicitly assumes that, if the genetic architecture is insensitive to sample characteristics, the genetic correlation is expected to be 1. However, this includes the expectation that trait-specific, study-specific sampling noise does not affect the expected genetic correlation. To test this hypothesis, we took random subsets of the UK Biobank, performed GWAS for various traits in each subset, and examined whether the genetic correlation is approximately one across traits, despite the traits having heterogeneous levels of sampling noise. In particular, we show pairwise genetic correlations across four datasets: (1) All of Us (AoU); (2) Two random subsets of the UKB (UKB1: 180,475 individuals; UKB2: 180,476 individuals); and GWAS summary statistics reported by the Neale lab who used a subsample of 361,194 unrelated European individuals (Neale). Points show estimates and vertical error bars show the 95% confidence intervals, as reported by bivariate `LDSC`. The dashed line marks a genetic correlation of 1. Alzheimer's disease (AD) is omitted from the above panels because the SNP heritability estimates in UKB were negative, precluding valid comparisons of genetic correlations. See **Text S1** for complete methods.

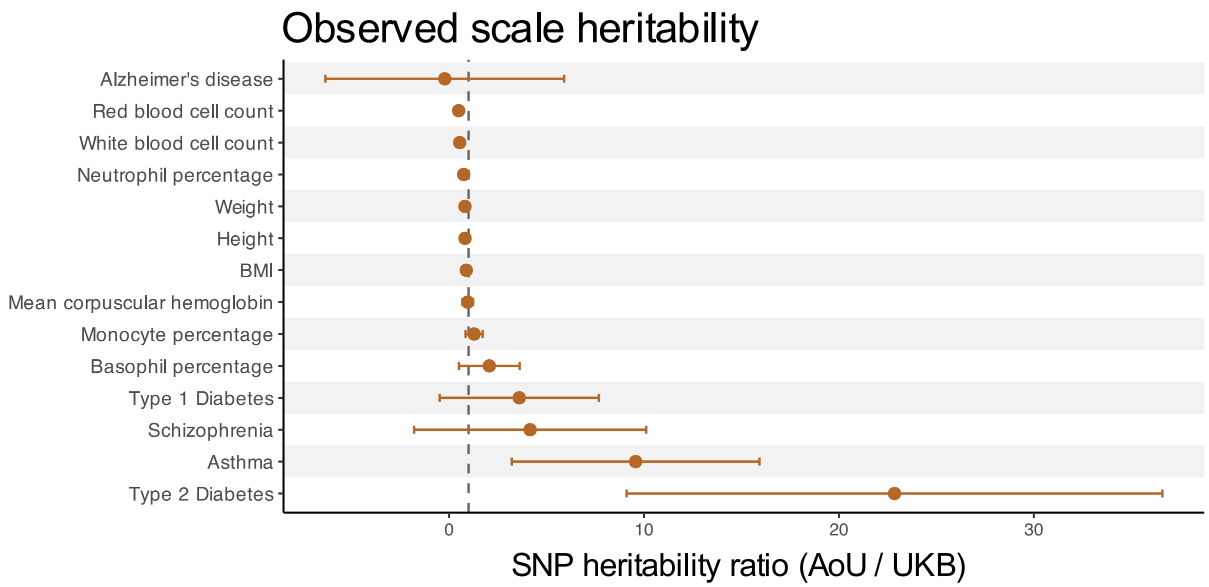

**Figure S10: Observed scale SNP heritability in UKB and AoU.** This figure mirrors the analysis of **Fig. 1A**, except the ratio of the observed scale SNP heritabilities for all traits are shown. Points represent estimates and horizontal bars represent 95% confidence intervals. The dashed vertical line corresponds to a ratio of 1.

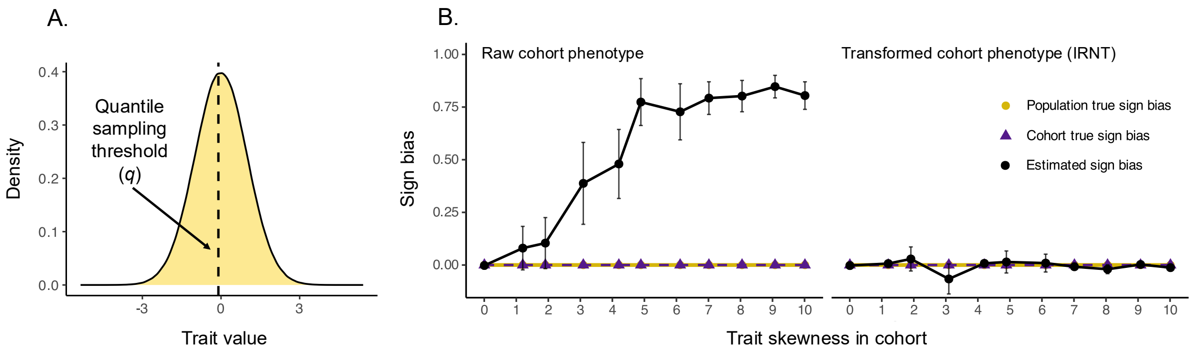

**Figure S11: Simulations demonstrate that cohort skewness induces sign bias when all SNP effects are null.** We sought to investigate whether cohort skewness could induce spurious genotype-phenotype associations leading to sign bias. **(A)** Following the procedures outlined under **Simulating traits and associated allelic effects** in the main text **Methods** for simulation scheme A, we generated a population of $2 \times 10^6$ individuals with a normally distributed trait and SNPs with no true effects on the trait in the population. Non-random cohorts were generated by sampling 10,000 individuals with varying trait skewness above a quantile threshold ($q$) of the trait distribution in the population. **(B)** For each realized skewness value in the cohorts, we performed GWAS either using the raw cohort trait distribution or the cohort trait distribution after inverse-rank normalization (IRNT). Points show the sign bias across 20 replicate cohorts, with error bars indicating 95% confidence intervals. Results are shown for variants with minor allele frequencies up to 1%. Since all SNPs have zero effect on the trait by design, there is no sign bias in the population (yellow) or in the cohort (dark purple). However, estimated sign bias (black) increased with increasing skewness in the cohort. IRNT eliminates the skewness of the analyzed trait and thus estimates of sign bias remained near zero. These results demonstrate that skewed cohorts generate directional asymmetry in the estimated sign of GWAS effects even when no variants affect the trait.
